# Robust and annotation-free analysis of alternative splicing across diverse cell types in mice

**DOI:** 10.1101/2021.04.27.441683

**Authors:** Gonzalo Benegas, Jonathan Fischer, Yun S. Song

## Abstract

Although alternative splicing is a fundamental and pervasive aspect of gene expression in higher eukaryotes, it is often omitted from single-cell studies due to quantification challenges inherent to commonly used short-read sequencing technologies. Here, we undertake the analysis of alternative splicing across numerous diverse murine cell types from two large-scale single-cell datasets—the *Tabula Muris* and BRAIN Initiative Cell Census Network—while accounting for understudied technical artifacts and unannotated isoforms. We find strong and general cell-type-specific alternative splicing, complementary to total gene expression but of similar discriminatory value, and identify a large volume of novel isoforms. We specifically highlight splicing variation across different cell types in primary motor cortex neurons, bone marrow B cells, and various epithelial cells; and show that the implicated transcripts include many genes which do not display total expression differences. To elucidate the regulation of alternative splicing, we build a custom predictive model based on splicing factor activity, recovering several known interactions while generating new hypotheses, including potential regulatory roles for novel alternative splicing events in critical genes including *Khdrbs3* and *Rbfox1*. We make our results available using public interactive browsers to spur further exploration by the community.

## Introduction

The past decade’s advances in single-cell genomics have enabled the data-driven characterization of a wide variety of distinct cell populations. Despite affecting more than 90% of human pre-mRNAs [1], isoform-level variation in gene expression has often been ignored because of quantification difficulties when using data from popular short-read sequencing technologies such as 10x Genomics Chromium and Smart-seq2 [2]. Long-read single-cell technologies, which greatly simplify isoform quantification, are improving [3, 4, 5, 6, 7], but remain more costly and lower-throughput than their short-read counterparts. For these reasons and others, short-read datasets predominate and we must work with short-reads to make use of the rich compendium of available data. In response, researchers have developed several computational methods to investigate splicing variation despite the sizable technical challenges inherent to this regime. A selection of these challenges and methods are summarized in Supplementary Text.

To complement single-cell gene expression atlases, we endeavored to analyze alternative splicing in large single-cell RNA-seq (scRNA-seq) datasets from the *Tabula Muris* consortium [8] and BRAIN Initiative Cell Census Network (BICCN) [9]. These data span a broad range of mouse tissues and remain largely unexplored at the isoform level. During our initial analyses, we encountered pervasive coverage biases, a heretofore largely unappreciated mode of technical variation which greatly confounds biological variation across cell types. Unsatisfied with the performance of current methods when confronted by these biases, we implemented our own suite of computational tools which allowed us to continue our analyses in a robust and annotation-free manner.

We find a strong signal of cell-type-specific alternative splicing and demonstrate that cell type can be accurately predicted given only splicing proportions. Moreover, our annotation-free approach enables us to detect a large quantity of cell-type-specific novel isoforms. In certain cell types, particularly the neuron subclasses, as many as 30% of differential splicing events that we detect are novel. In general, across the many considered cell types and tissues in both datasets, we find only a narrow overlap between the top differentially expressed and the top differentially spliced genes within a given cell type, illustrating the complementarity of splicing to expression. Our examination of neurons in the primary motor cortex suggests that splicing distinguishes neuron classes and subclasses as readily as does expression. We showcase alternative splicing patterns specific to the GABAergic (inhibitory) and Glutamatergic (excitatory) neuron classes as well as the subclasses therein. The implicated transcripts include key synaptic molecules and genes which do not display expression differences across subclasses. In developing marrow B cells, we find alternative splicing and novel transcription start sites (TSS) in critical transcription factors such as *Smarca4* and *Foxp1*, while further investigation reveals dissimilar trajectories for expression and alternative splicing in numerous genes across B cell developmental stages. These findings buttress our belief in the complementary nature of these processes and provide clues to the regulatory architecture controlling the early B cell life cycle. To facilitate easy exploration of these datasets and our results, we make available several interactive browsers as a resource for the genomics community.

Finally, to advance our understanding of alternative splicing regulation, we build a statistical machine learning model to predict splicing events by leveraging both the expression levels and splicing patterns of splicing factors across cell types. This model recovers several known regulatory interactions such as the repression of splice site 4 exons in neurexins by *Khdrbs3*, while generating new hypotheses for experimental follow-up. For example, in addition to the regulatory effect of the whole-gene *Khdrbs3* expression, the model predicts a regulatory role for a novel alternative TSS in this gene. In aggregate, our results imply that alternative splicing serves as a complementary rather than redundant component of transcriptional regulation and supports the mining of large-scale single-cell transcriptomic data via careful modeling to generate hypothetical regulatory roles for splicing events.

## Results

### Methods overview

#### Robust, annotation-free quantification based on alternative introns

Most methods rely on the assumption that coverage depth across a transcript is essentially uniform (e.g., *Akr1r1*, Figure S1a). We instead found that Smart-seq2 data [2] frequently contain sizable fractions of genes with coverage that decays with increasing distance from the 3’ ends of transcripts. For example, in mammary gland basal cells from the *Tabula Muris* data set [8], *Ctnbb1* shows a gradual drop in coverage (Figure S1b) while *Pdpn* displays an abrupt reduction halfway through the 3’ UTR (Figure S1c). That the magnitude of these effects varies across technical replicates (plates) suggests they could be artifacts, possibly related to degradation or interrupted reverse transcription. Similar coverage bias artifacts are also apparent in the BICCN primary motor cortex data [9] (Figure S2).

Such coverage biases affect gene expression quantification, and in some cases these batch effects are sufficient to comprise a significant proportion of the observed variation in expression levels. For the *Tabula Muris* mammary gland data set, a low-dimensional embedding of cells based on gene expression reveals that some cell type clusters exhibit internal stratification by plate (Figure 1a). A subsequent test of differential gene expression between plate B002438 and all other plates returns 2,870 significant hits after correction for multiple hypothesis testing, and all manually inspected differentially expressed genes exhibit these types of coverage biases. Perhaps unsurprisingly, quantification at the transcript level is apt to be even more sensitive to these artifacts than gene-level quantification, especially if it is based on coverage differences across the whole length of the transcript (denoted as *global* isoform quantification in Table S1). The UMAP embedding of isoform proportions (kallisto [10]), exon proportions (DEXSeq [11]), 100 bp bin coverage proportions (ODEGR-NMF [12]) or junction usage proportions across the whole gene (DESJ [13]) depicts a stronger plate clustering pattern which supersedes the anticipated cell type clusters (Figure 1b-e).

**Figure 1:**
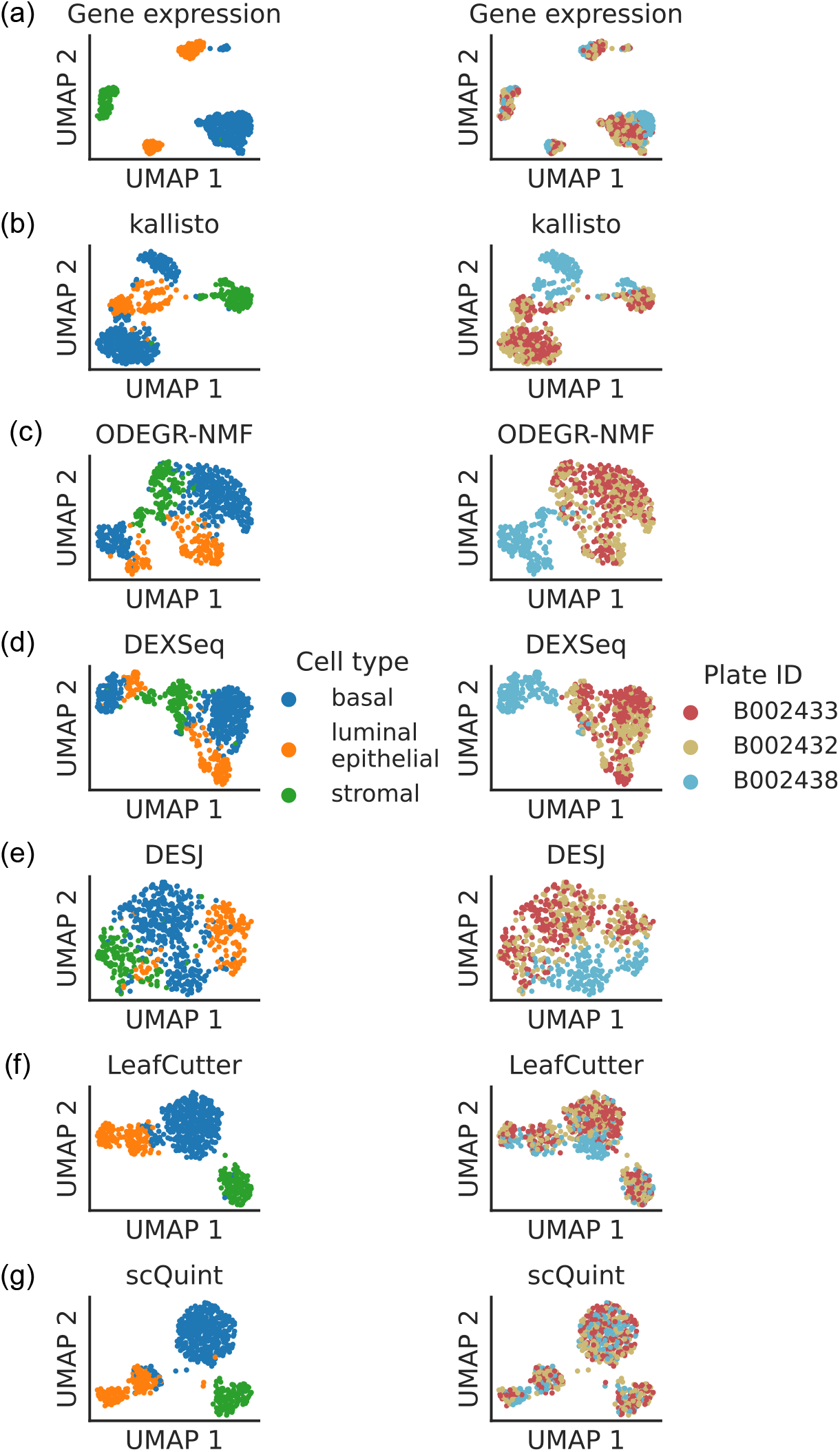
Clustering patterns by cell type and plate in the mammary gland from a three month-old female mouse in Tabula Muris. Cell embeddings based on different features were obtained by running PCA (gene expression) or VAE (the rest) followed by UMAP and subsequently colored by cell type (left column) and the plate in which they were processed (right column). (a) Gene expression, quantified using featureCounts (log-transformed normalized counts). (b) Isoform proportions. Isoform expression was estimated with kallisto and divided by the total expression of the corresponding gene to obtain isoform proportions. (c) Coverage proportions of 100 base-pair bins along the gene, as proposed by ODEGR-NMF. (d) Exon proportions, as proposed by DEXSeq. (e) Intron proportions across the whole gene, as proposed by DESJ. (f) Alternative intron proportions quantified by LeafCutter. (g) Alternative intron proportions (for introns sharing a 3’ acceptor site) as quantified by scQuint.

With these considerations in mind, we sought to quantify isoform variation in a local manner that would be more robust to coverage differences along the transcript. One such approach is alternative intron quantification, as performed by bulk RNA-seq methods MAJIQ [14], JUM [15] and LeafCutter [16]. Promisingly, quantification via LeafCutter (Figure 1f) yields an embedding that displays less clustering by plate than global quantifications such as kallisto or DESJ. Inspired by LeafCutter, our method scQuint (<u>s</u>ingle-<u>c</u>ell <u>qu</u>antification of <u>int</u>rons) also quantifies alternative introns but restricts to those sharing a common 3’ acceptor site (Figure 2). As a result, alternative events are equidistant from the 3’ end of transcripts and are not affected by the coverage biases described above. The embedding of cells based on our quantification approach (Figure 1g) shows less clustering by plate than LeafCutter and other methods. Slight clustering by plate can still be seen in luminal epithelial cells, which could be caused by other kinds of technical artifacts such as sequence bias. As we describe later, scQuint also provides more accurate differential splicing *p*-values compared to LeafCutter.

**Figure 2:**
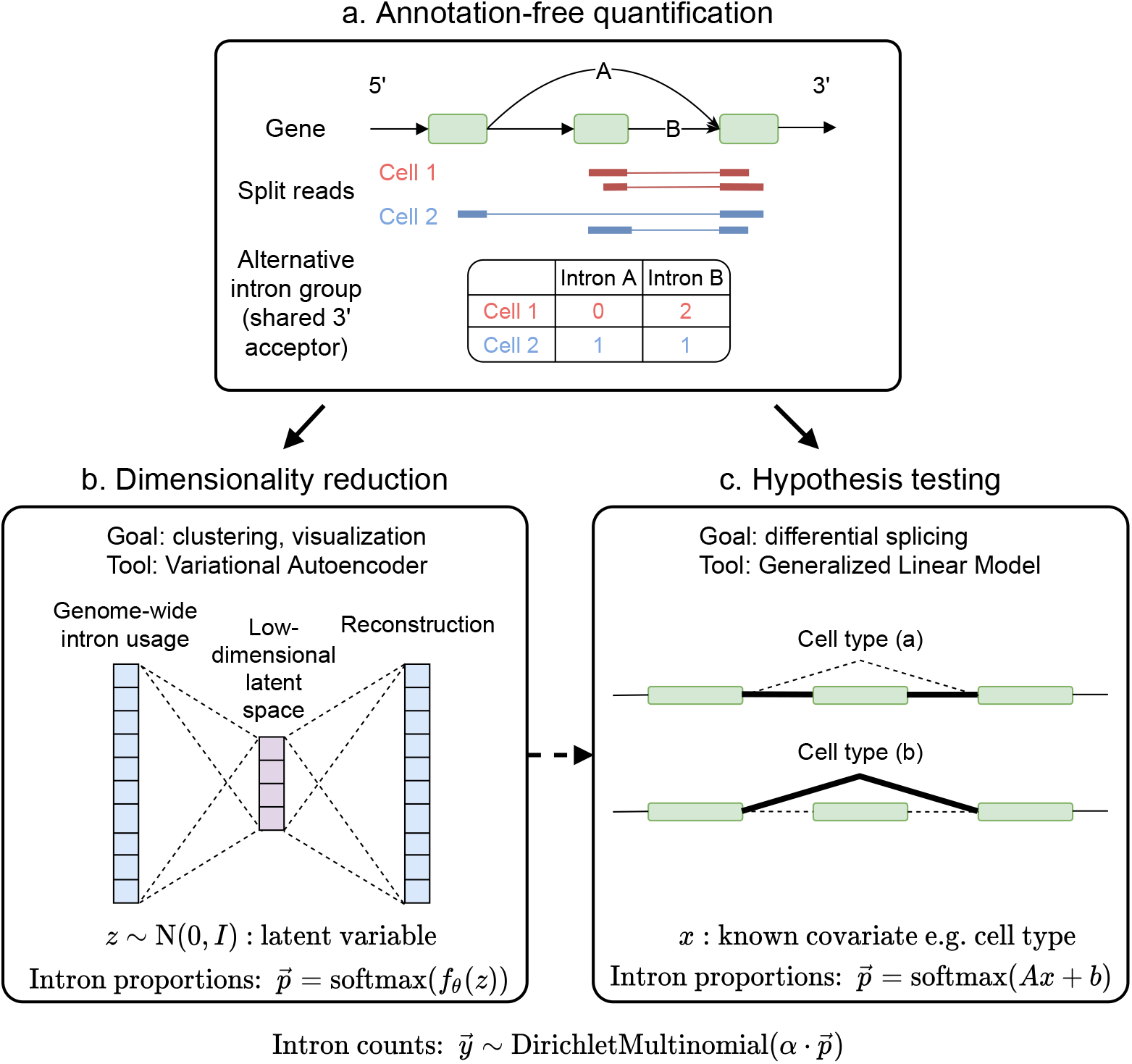
Overview of scQuint. (a) Intron usage is quantified from split reads in each cell, with introns sharing 3’ splice sites forming alternative intron groups. (b) Genome-wide intron usage is mapped into a low dimensional latent space using a Dirichlet-Multinomial VAE. Visualization of the latent space is done via UMAP. (c) A Dirichlet-Multinomial GLM tests for differential splicing across conditions such as predefined cell types or clusters identified from the splicing latent space.

Another advantage of alternative intron quantification is the ability to easily discover novel isoforms. Whereas short reads generally cannot be associated with specific transcript isoforms, nor even exons, if they partially overlap, split reads uniquely associate with a particular intron. Consequently, intron-based quantification does not depend on annotated transcriptome references and allows for novel isoform discovery. This is important since, as detailed later, we estimate up to 30% of novel cell-type-specific differential splicing events. Expedition [17] and ASCOT [18] also perform annotation-free quantification based on local intron usage, but they consider only simple binary events and ignore other important events such as alternative transcription start sites, which we find to be plentiful in our analysis. Moreover, neither Expedition nor ASCOT provide a statistical test for differential splicing across cell subpopulations.

#### Dimensionality reduction with Variational Autoencoder

To perform dimensionality reduction using splicing profiles, we developed a novel Variational Autoencoder (VAE) [19] with a Dirichlet-Multinomial noise model (Figure 2b, Materials and Methods). VAEs are flexible and scalable generative models which have been successfully applied to analyze gene expression [20] but have not yet been employed to investigate alternative splicing. We compared the latent space obtained with the VAE to the one obtained using Principal Component Analysis (PCA), a standard dimensionality reduction technique used in the LeafCutter software package. The VAE is able to better distinguish cell types than can PCA (Figure 3). While cortex cells are already quite well separated in the PCA latent space, the increased resolution offered by the VAE is better appreciated in other tissues such as mammary gland and diaphragm.

**Figure 3:**
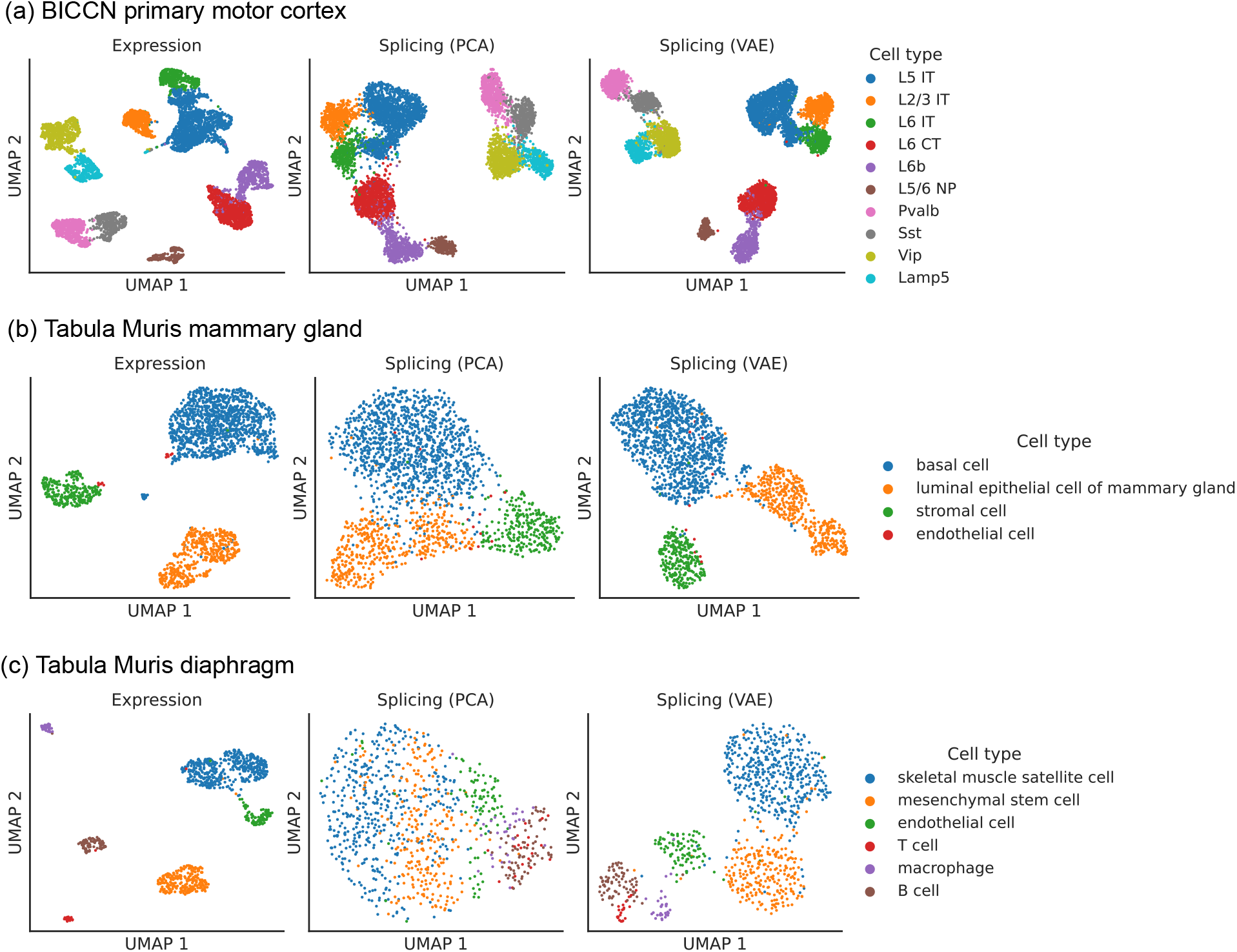
Comparison of splicing latent spaces obtained with PCA and VAE. Cells from (a) the cortex, (b) mammary gland and (c) diaphragm are projected into a latent space using PCA or VAE and visualized using UMAP. Cell type labels are obtained from the original data sources and are based on clustering in the expression latent space. The VAE is able to better distinguish cell types in the splicing latent space than PCA.

#### Differential splicing hypothesis testing with Generalized Linear Model

To test for differential splicing across cell types or conditions, we adopt a Dirichlet-Multinomial Generalized Linear Model (GLM) coupled with a likelihood-ratio test (Figure 2c, Materials and Methods). We leverage one of the models proposed by LeafCutter for bulk RNA-seq, apply it to our Smart-seq2 intron quantification, and find that it yields well-calibrated *p*-values (Figure S3). LeafCutter’s differential splicing p-values, however, are not as well calibrated (Figure S3), which could be due to differences in the model or its optimization. Since LeafCutter’s *p*-values are anti-conservative, note that it can lead to more false positives than expected.

### Augmenting cell atlases with splicing information

We applied scQuint to two of the largest available Smart-Seq2 data sets. The first comprehensively surveys the mouse primary motor cortex (*BICCN Cortex*) [9] while the second contains over 100 cell-types distributed across 20 mouse organs (*Tabula Muris*) [8] (Table S2). We detect more alternative introns in *BICCN Cortex* neurons than in the entire broad range of cell types present in *Tabula Muris* (which includes neurons but in much smaller number). This observation comports with previous findings that the mammalian brain has exceptionally high levels of alternative splicing [21]. Booeshagi et al. [22] analyzed *BICCN Cortex* at the transcript level, but focused on changes in absolute transcript expression rather than proportions. While the authors indirectly find some differences in transcript proportions by inspecting genes with no differential expression, this is not a systematic analysis of differential transcript usage. Meanwhile, only microglial cells in *Tabula Muris* [23] have been analyzed at the isoform level. (*Tabula Muris* also contains 10x Chromium data analyzed at the isoform level [24]).

As a community resource, we provide complementary ways to interactively explore splicing patterns present in these data sets (Figure 4), available at https://github.com/songlab-cal/scquint-analysis/ with an accompanying tutorial video. The UCSC Genome Browser [25] permits exploration of alternative splicing events within genomic contexts such as amino acid sequence, conservation score, or protein binding sites, while allowing users to select different length scales for examination. We additionally leverage the cell × gene browser [26] (designed for gene expression analysis) to visualize alternative intron PSI (percent spliced-in, defined as the proportion of reads supporting an intron relative to the total in the intron group) via cell embeddings. Further, one can generate histograms to compare across different groups defined by cell type, gender, or even manually selected groups of cells. These tools remain under active development by the community, and we hope that both the genome- and cell-centric views will soon be integrated into one browser.

**Figure 4:**
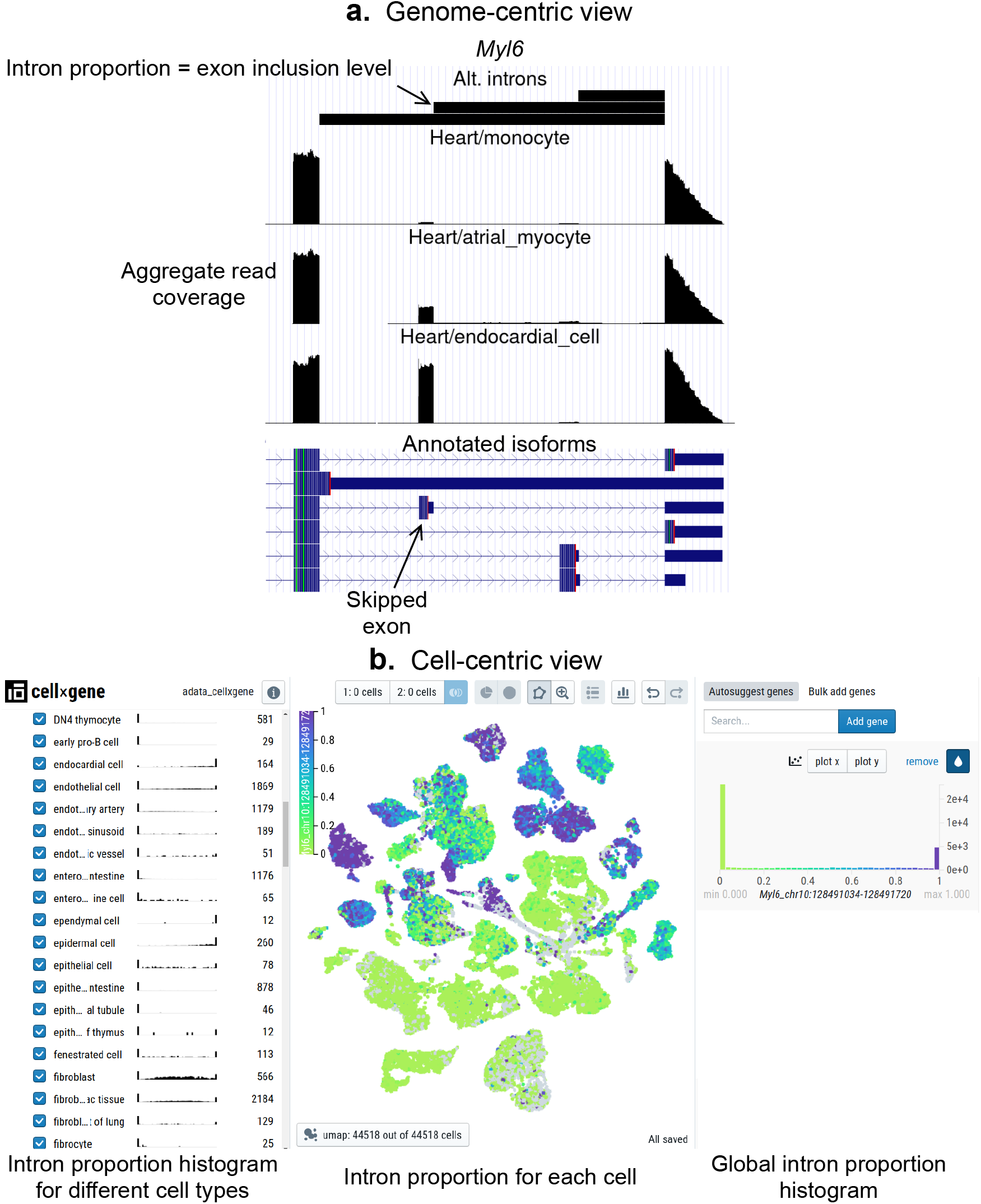
Interactive visualizations of splicing patterns. As an example, a skipped exon in *Myl6*. (a) The UCSC Genome browser visualization of this locus. Bottom: annotated isoforms of *Myl6*, including a skipped exon. Center: aggregate read coverage in three cell types with varying inclusion levels of the skipped exon. Top: three alternative introns that share a 3’ acceptor site. The identified intron’s proportion corresponds to the skipped exon’s inclusion level. (b) cell×gene browser visualization of the marked intron’s proportions (`Myl6_chr10:128491034-128491720`). Center: intron proportion for each cell in the UMAP expression embedding. Sides: intron proportion histogram for (left) different cell types and (right) all cells.

### Cell-type-specific splicing signal is strong and complementary to gene expression

#### Primary motor cortex

We first explored the splicing latent space of *BICCN Cortex* cells by comparing it to the usual expression latent space (Figure 5a). Cells in the splicing latent space strongly cluster by cell type (annotated by Yao *et al*. [9] based on gene expression). A similar analysis was recently performed [27] on a different cortex subregion in which most, but not all, neuron subclasses could be distinguished based on splicing profiles (e.g., L6 CT and L6b could not be separated). However, the authors only considered annotated skipped exons, a subset of the events we quantify, and used a different dimensionality reduction technique.

**Figure 5:**
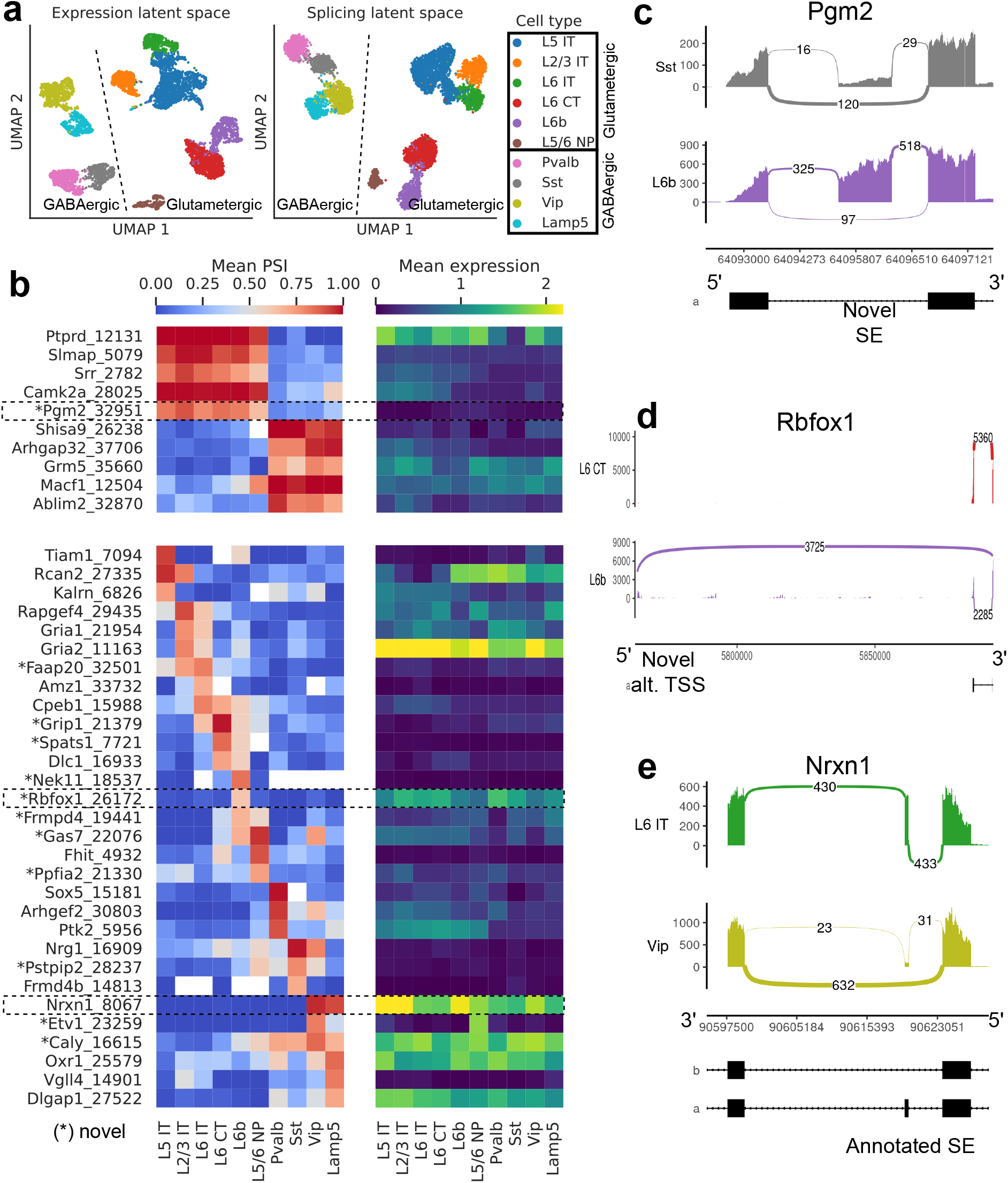
Splicing patterns in *BICCN Cortex*. (a) Expression and splicing latent spaces, visualized using UMAP. The expression (splicing) latent space is defined by running PCA (VAE) on the gene expression (alternative intron proportion, PSI) matrix. Cell types separate well in both latent spaces. (b) PSI of selected introns (left) and expression (log-transformed normalized counts) of their respective genes (right) averaged across cell types. Top: introns distinguishing Glutamatergic and GABAergic neuron classes. Bottom: introns distinguishing neuron subclasses. Intron coordinates are available in Table S3. (c-e) Sashimi plots [32] of specific alternative splicing events, displaying overall read coverage with arcs indicating usage of different introns (certain introns are shrunk for better visualization). (c) Novel skipped exon in *Pgm2*. (d) Novel alternative transcription start site (TSS) in *Rbfox1*. (e) Annotated skipped exon (SE) in *Nrxn1*.

Figure 5b (top left) highlights some differentially spliced genes between Glutamatergic and GABAergic neurons, including the glutamate metabotropic receptor *Grm5* as well as *Shisa9/Ckamp44*, which associates with AMPA ionotropic glutamate receptors [28]. The expression pattern of these genes, meanwhile, does not readily distinguish the neuron classes (Figure 5b, top right). In *Pgm2*, a gene of the glycolysis pathway thought to be regulated in the developing cortex by mTOR [29], we discover a novel exon preferentially included in Glutamatergic neurons (Figure 5c).

Our differential splicing test reveals thousands of cell-type-specific splicing events, highlighting marker introns that distinguish neuron subclasses, while the expression of their respective genes does not; e.g., compare bottom left and bottom right panels of Figure 5b. As another example of the many novel isoforms we discover, we showcase a novel alternative transcription start site in *Rbfox1*, a splicing factor known to regulate cell-type-specific alternative splicing in the brain [30] (Figure 5d). This novel TSS (`Rbfox1_26172, chr16:5763871-5763913`), which lies in a highly-conserved region, is (partially) used by only L6b neurons. We are also able to detect well-known cell-type-specific alternatively spliced genes such as *Nrxn1*, which encodes a key pre-synaptic molecule (Figure 5e) [31]. In this case, we observe an exon (known as splice site 2) exclusively skipped in Vip and Lamp5 neurons.

#### General patterns in *Tabula Muris*

We next turned our attention to *Tabula Muris*, which comprises a wide variety of organs and cell types from across the entire body. As before, we initially compared the expression and splicing latent spaces using UMAP (Figure 6a). This revealed broadly consistent clusters between projections, but a visible shift in the global layout of these clusters. In particular, whereas cell types were better separated in the expression projection, cell classes (e.g., endothelial, epithelial, immune) formed more coherent clusters in the splicing projection.

**Figure 6:**
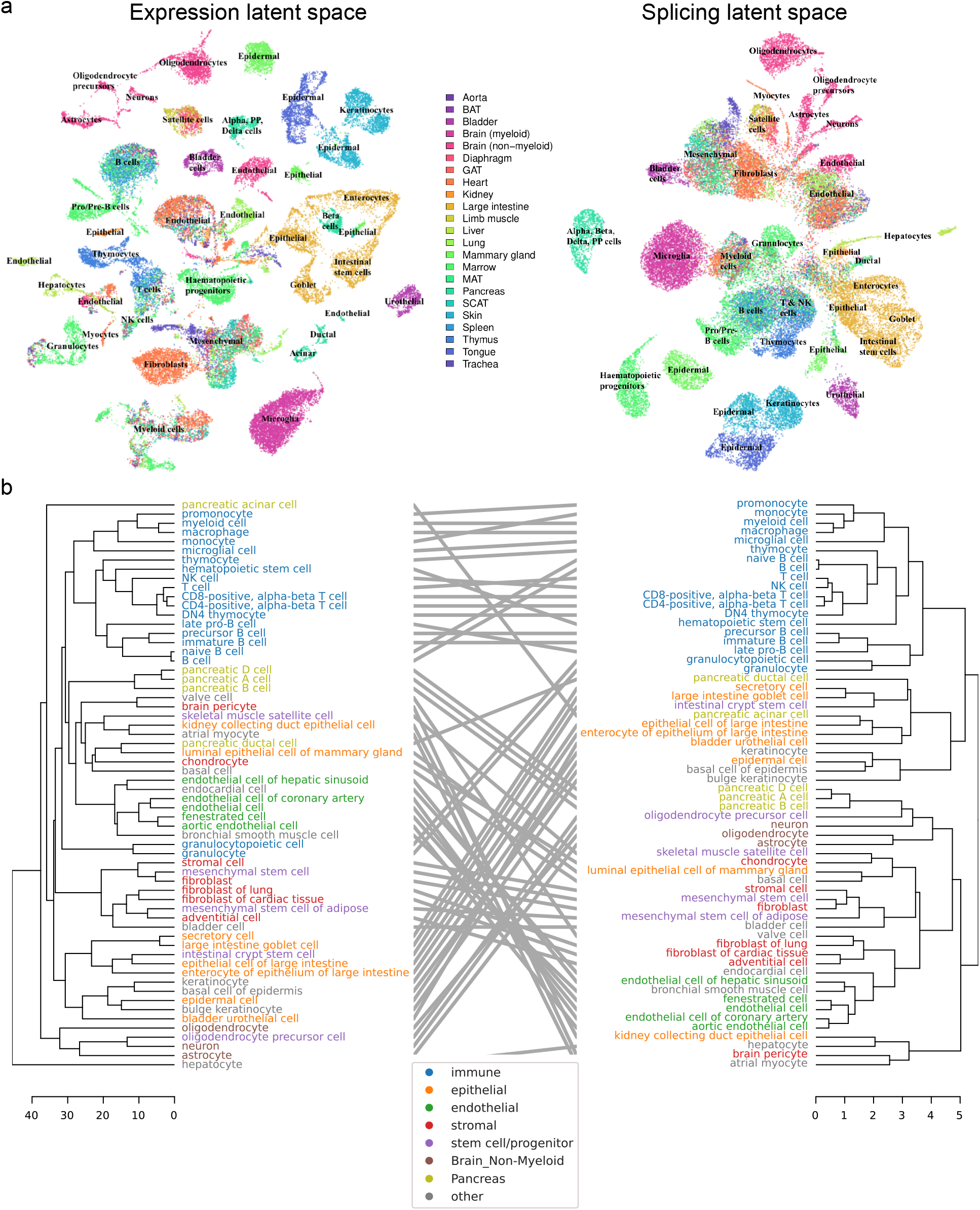
Global analysis of *Tabula Muris*. (a) UMAP visualization of the expression (left) and splicing (right) latent spaces. Each dot is a cell, colored by organ, and overlays indicate the primary cell type comprising that cluster. (b) Tanglegram comparing dendrograms of major cell types based on distances in the expression (left) and splicing (right) latent spaces, highlighting functional classes with specific colors.

To supplement our qualitative comparison of UMAP projections with a more rigorous approach, we built dendrograms and a tanglegram using the respective distances between cells in each of the expression and splicing latent spaces (Figure 6b). Despite minor shifts, the dendrograms resemble one another, and most subtree structure is preserved. The low value of their entanglement, a quantitative measure of the discrepancy between hierarchical clusterings, at only 6% indicates a high degree of similarity. (For comparison, the entanglement value between the dendrogram for all expressed genes and that for transcript factors is 11% [8].) As in the UMAP visualization, immune cells group together more closely in the splicing dendrogram. However, unlike the UMAP projection, we observe that several types of pancreatic cells cluster together with neurons, a cell type long believed to share an evolutionary origin [33]. Notably, the left dendrogram in Figure 6b shows that hepatocytes are clear outliers in the expression latent space. We suspect this may be due to technical differences from using 96-well plates rather than the 384-well plates used for other cell types.

#### B cell development in the marrow

We then focused on developing B cells from the bone marrow in *Tabula Muris*. In the splicing latent space, we found that immature B cells are harder to distinguish from the other B cell subpopulations (Figure 7a), reflecting less refined splicing programs or limitations in transcript capture efficiency. The several differential splicing events we identified throughout development displayed splicing trajectories mostly independent from the trajectories of gene expression (Figure 7b). We highlight alternative TSSs (one of them novel) in two transcription factors essential for B cell development: *Smarca4*, encoding BRG1 [34] (Figure 7c); and *Foxp1* [35] (Figure 7d). While *Foxp1* expression peaks in pre-B cells and does not follow a monotonic trend over developmental stages, the alternative TSS is progressively included throughout B cell development. Combining gene-level expression with TSS usage, which can influence translation rate, provides a more nuanced characterization of the expression patterns of these important transcription factors. Some other differentially spliced genes with well-known roles in B cell development are *Syk* [36], *Dock10* [37], *Selplg/Psgl-1* [38], and *Rps6ka1* [39].

**Figure 7:**
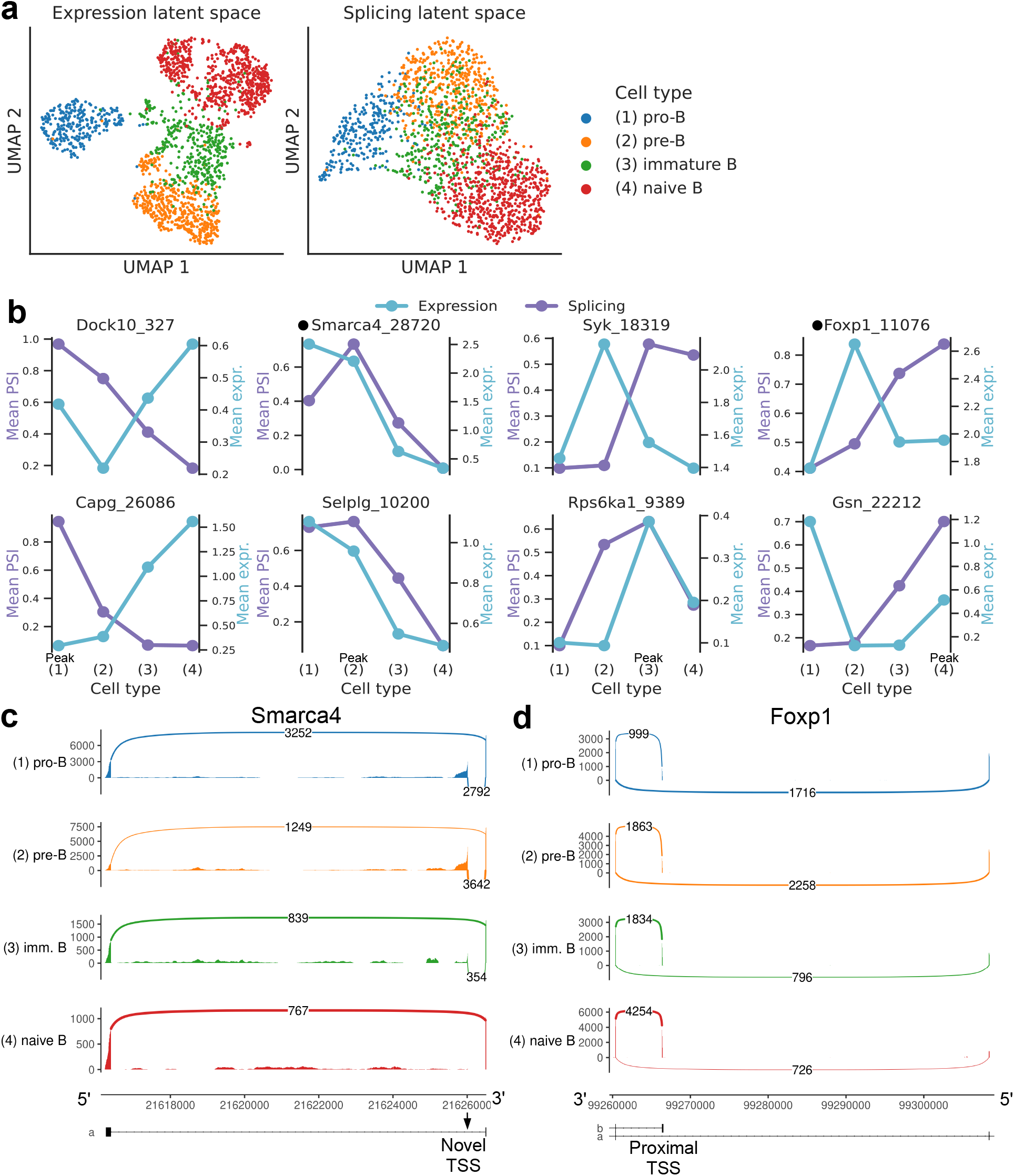
Splicing in developing marrow B cells from *Tabula Muris*. B cell developmental stages include pro-B, pre-B, immature B, and naive B. (a) Expression versus splicing latent space, as defined previously. In the splicing latent space, some cells types (pro-B) are better distinguished than others (immature B). (b) PSI of some introns that are differentially spliced throughout development, together with expression of the respective genes (log-transformed normalized counts). Expression and splicing can have very different trajectories. Intron coordinates are available in Table S4. (c) Sashimi plot of novel alternative transcription start site (TSS) in *Smarca4*. The novel TSS has maximum usage in pre-B cells, and then decays, while the expression peaks at pro-B cells. (d) Sashimi plot of an annotated alternative TSS in *Foxp1*. The proximal TSS in increasingly used as development progresses, while the expression peaks at pre-B cells.

#### Epithelial and endothelial cell types across organs

Having compared different cell types within organs, we analyzed putatively similar cell types which are present in multiple organs to investigate splicing variation associated with tissue environment and function. We find many alternative introns with strong PSI differences across epithelial cell types, including several which are novel (Figure 8a). Conversely, apart from those in the brain, endothelial cell types fail to display such striking differences (Figure 8b). These patterns are consistent with the UMAP projection and dendrogram, both of which suggested less heterogeneity among endothelial than epithelial cells (Figure 6).

**Figure 8:**
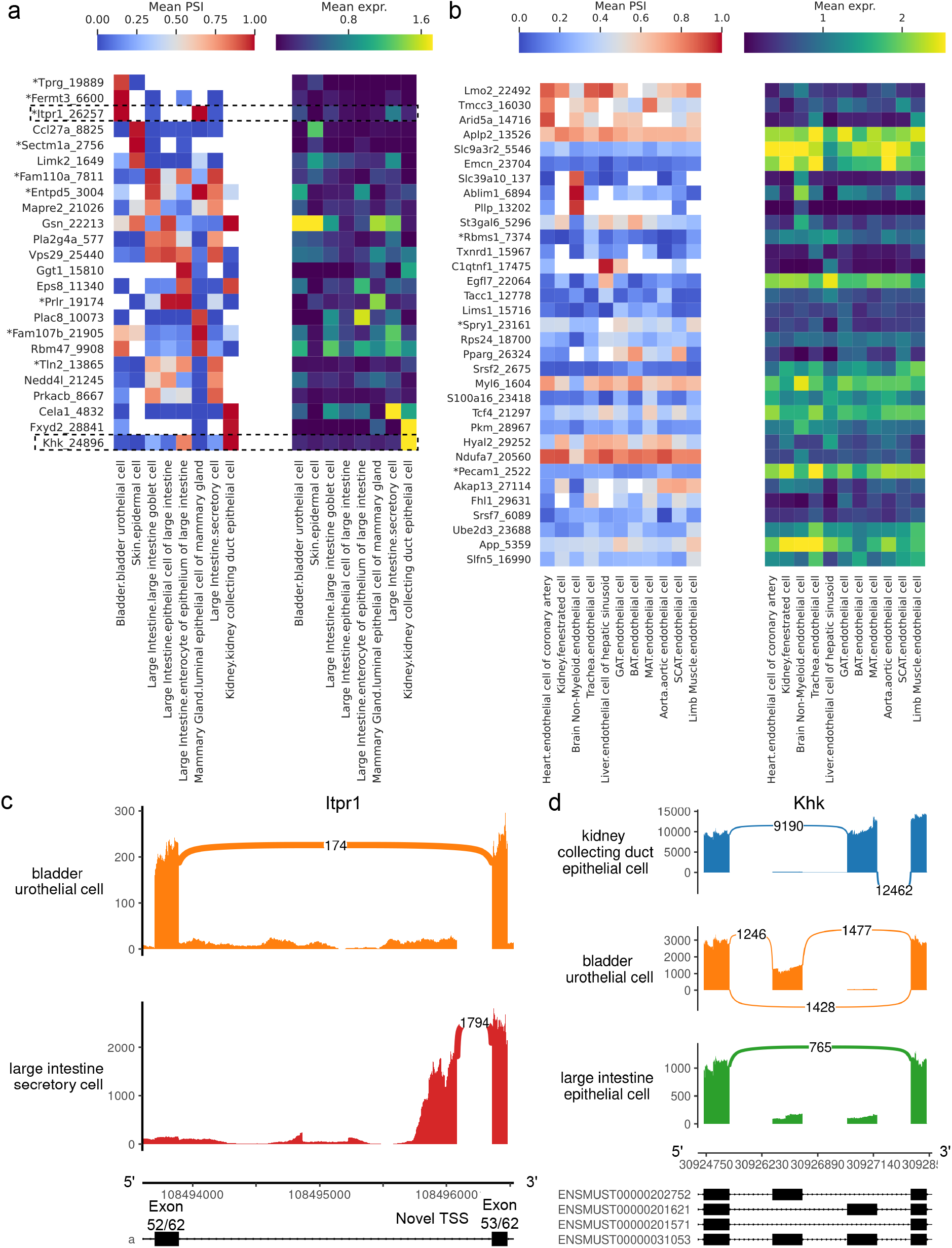
Alternative splicing patterns across epithelial and endothelial cell types. (a-b) PSI of selected introns (left) and expression (log-transformed normalized counts) of the corresponding genes (right) averaged across cell types. Novel intron groups are marked with (*). (a) Introns distinguishing epithelial cell types. (b) Introns distinguishing endothelial cell types. Intron coordinates are available in Table S5. (c) Sashimi plot of an alternative TSS in *Itpr1* (full-gene view in Figure S4). (d) Sashimi plot of a complex alternative splicing event in *Khk*.

Our analysis revealed a novel alternative TSS in *Itpr1* (Figure 8c), an intracellular calcium channel in the endoplasmic reticulum, which regulates secretory activity in epithelial cells of the gastrointestinal tract [40]. This novel TSS yields a shorter protein isoform (full view in Figure S4) which preserves the transmembrane domain, though it is unclear whether this isoform is functional. Notably, it is the predominant isoform in large intestine secretory cells, and these cells express *Itpr1* at the highest level among all epithelial cell types in the dataset. All nine novel alternative splicing events in Figure 8a are alternative TSSs, with four affecting the 5’ UTR and five affecting the coding sequence.

Figure 8d illustrates a complex alternative splicing event in *Khk* involving the well-studied exons 3a and 3c [41]. Khk catalyzes the conversion of fructose into fructose-1-phosphate, and the two protein isoforms corresponding to either exon 3a or 3c inclusion differ in their thermostability and substrate affinity [42]. While the literature describes these exons as mutually exclusive, the transcriptome reference includes transcripts where neither or both may be included. Although we did not find cell types with high inclusion rates for both exons, we did see multiple cell types where both exons are predominantly excluded, e.g., epithelial cells from the large intestine. Other differentially spliced genes are involved in cellular junctions, which are particularly important in epithelial tissue. These include *Gsn*, *Eps8*, *Tln2*, *Fermt3*, and *Mapre2*.

#### Comparison of selected tissues

Because of the breadth of the *Tabula Muris* data set, we can look for general trends across a diverse array of tissues and cell types. Table 1 summarizes differential expression and splicing for some of the cell types and tissues with the largest sample sizes. First, we note the intersection between the top 100 most differentially expressed and top 100 most differentially spliced genes (ranked by *p*-value) is consistently low. This means that most differentially spliced genes, which might be of critical importance in a biological system, will go unnoticed if a study only considers differential expression. Second, L5 IT neurons have a larger fraction of genes with differential splicing relative to the number of differentially expressed genes.

**Table 1:** Summary of differential expression and splicing for select cell types with the largest sample sizes. The overlap between the top 100 differentially expressed genes and the top 100 differentially spliced genes is low, indicating that splicing provides complementary information. In addition, L5 IT neurons have a higher ratio of differentially spliced genes to differentially expressed genes than the other cell types. *Diff. spl. genes*: number of differentially spliced genes between the cell type and other cell types in the same tissue. *Diff. exp. genes*: number of differentially expressed genes between the cell type and other cell types in the same tissue. See Materials and Methods for details on the tests for differential splicing and expression.

| Tissue | Total<br># cells | # cell<br>types | Cell type | # cells | Diff.<br>spl.<br>genes | Diff.<br>exp.<br>genes | Ratio | Top-100<br>overlap |
| --- | --- | --- | --- | --- | --- | --- | --- | --- |
| Brain Non-Myeloid | 3049 | 6 | Oligodendrocyte | 1390 | 880 | 8835 | 0.10 | 4 |
| Cortex | 6220 | 10 | L5 IT | 1571 | 1447 | 6402 | 0.23 | 2 |
| Heart | 4144 | 6 | Endothelial cell of coronary artery | 1126 | 465 | 7108 | 0.07 | 5 |
| Large Intestine | 3729 | 5 | Enterocyte of epithelium | 1112 | 586 | 10786 | 0.05 | 2 |
| Marrow | 4783 | 10 | Hematopoietic stem cell | 1363 | 692 | 9909 | 0.07 | 2 |

We found many more cell-type-specific differential splicing events in the cortex than in the marrow, as well as a higher proportion of events involving novel junctions, which can reach 30% (Figure 9a). Most differential splicing events that we detected with alternative introns fall in the coding portion of the gene, with high proportions in the 5’ UTR (Figure 9b). We find an increased proportion of differentially spliced non-coding RNA in the cortex, the majority of which are previously unannotated events. To systematically evaluate how well cell types can be distinguished in the expression and splicing latent spaces, we calculated the ROC AUC score for the one-versus-all classification task for each cell type in each tissue using a binary logistic regression model (Figure 9c). Since cell type labels were defined using gene expression values, near-perfect classification is to be expected using the expression latent space. Classification based only on the splicing latent space is very good in general, suggesting that cell-type-specific differential splicing is rather pervasive. A few cell types were more challenging to classify correctly using splicing patterns alone. One such example is immature B cells, a reflection of the lower degree of separation observed in the embedding of Figure 7a.

**Figure 9:**
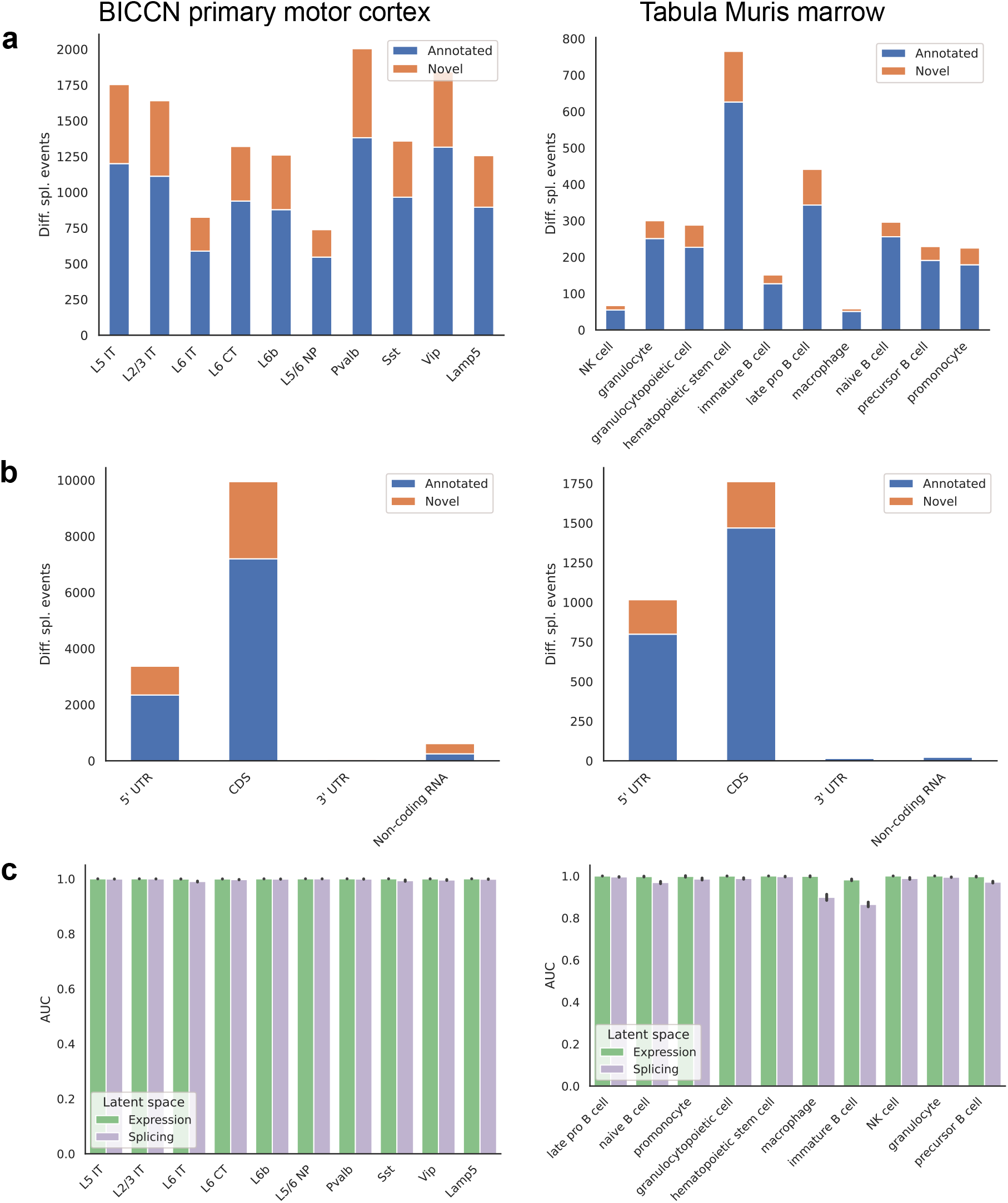
Patterns across tissues. (a) Number of differential splicing events detected in each cell type. Cortex cell types have more differential splicing events and larger proportions of novel events (those involving an intron absent from the reference). (b) Number of differential splicing events in different gene regions aggregated over cell types. Cortex cell types have higher proportions of events in coding regions and noncoding RNAs. Note: y-axes are not on the same scale. (c) ROC AUC score for classification of each cell type versus the rest based on either the expression or splicing latent space, using logistic regression. The score for splicing-based classification is near-perfect in most cell types with some exceptions such as immature B cells in the marrow.

### Finding splicing factors associated with specific alternative splicing events

Several splicing factors have been identified as regulators of specific alternative splicing events, but most regulatory interactions remain unknown (see [43] for a review focused on the brain). To complement expensive and laborious knockout experiments, we sought to generate regulatory hypotheses by analyzing the correlation between splicing outcomes and splicing factor variation across cell types. Focusing on a subset of highly expressed genes in BICCN primary motor cortex neurons, we fit a sparse linear model regressing PSI of skipped exons on both expression and splicing patterns of splicing factors (Figure 10a and Figure S5). Our model recovers several known regulatory interactions such as Khdrbs3’s repression of splice site 4 (SS4) in neurexins, modulating their binding with post-synaptic partners [43]. Additionally, the proportion of a novel alternative TSS (though annotated in the human reference) in *Khdrbs3* (Figure 10b) is negatively associated with SS4 in *Nrxn1* and *Nrxn3*. The skipping of exon 5 (E5) of *Grin1*, which controls long-term synaptic potentiation and learning [44], is known to be regulated by Mbnl2 and Rbfox1 [43]. The model associates *Grin1* E5 PSI with the expression of *Rbfox1* but not *Mbnl2*; however, it does suggest an association with the PSI of two skipped exons in *Mbnl2* (Figure 10c) and further implicates the inclusion level of the novel alternative TSS in *Rbfox1* reported above (`Rbfoxl_26172, chr16:5763912-6173605`, Figure 5d). These results help clarify the disparate impacts of expression and alternative splicing in splicing factors, and encourage the use of regression models to suggest candidate regulators of cell-type-specific alternative splicing. Such computationally generated hypotheses are particularly valuable for splicing events in splicing factors because of the heightened difficulty to experimentally perturb specific exons rather than whole genes.

**Figure 10:**
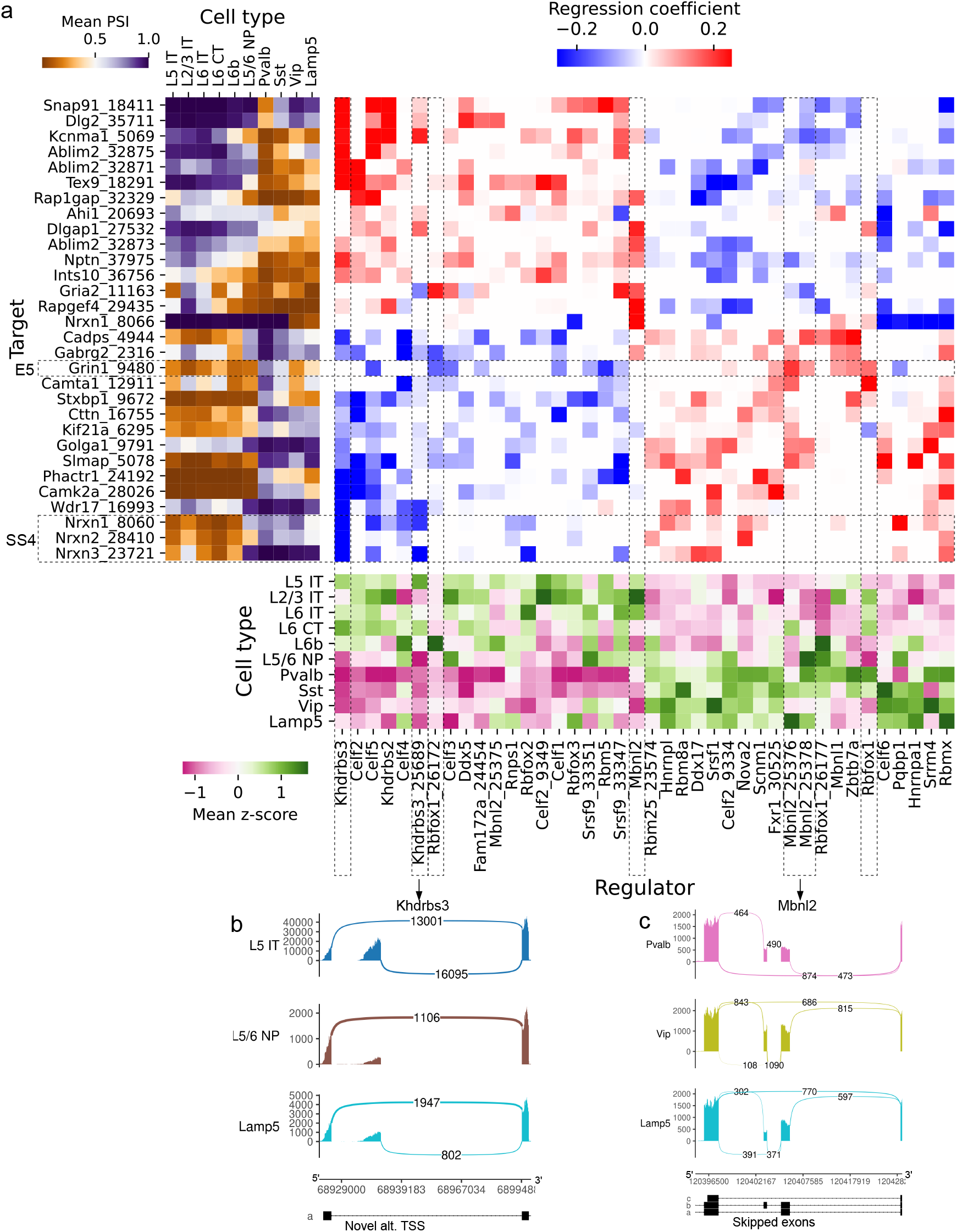
Associations between splicing factors and alternative splicing. (a) Regression analysis of exon skipping based on expression and splicing of splicing factors, using the BICCN mouse primary motor cortex dataset. Left panel: mean PSI of skipped exons across cell types. Bottom panel: mean z-scores of selected splicing factor features across cell types, including whole-gene expression (gene name) and PSI of alternative introns (gene name and numerical identifier). Intron coordinates are available in Table S6. Center panel: regression coefficients (log-odds) of each splicing factor feature used to predict skipped exon PSI in our sparse Dirichlet-Multinomial linear model. Full results are available in Figure S5. (b) Novel alternative TSS in *Khdrbs3*. (c) Annotated skipped exons in *Mbnl2*.

## Discussion

In this study, we successfully extend the analysis of two single-cell atlases to the level of alternative splicing, overcoming the usual technical challenges as well as coverage artifacts and incomplete isoform annotations. Our results indicate the presence of strong cell-type-specific alternative splicing and previously unannotated splicing events across a broad array of cell types. In most cases, splicing variation is able to differentiate cell types just as well as expression levels. We also note a striking lack of overlap between the most strongly differentially expressed and spliced genes (Table 1), suggesting that expression and splicing are complementary rather than integrated processes. Moreover, this complementarity may also manifest temporally, as we show in developing B cells in the marrow. Another outstanding question is the functional significance of isoforms, and we find that most differential splice sites appear in the coding sequence with a sizeable minority also mapping to 5’ UTRs. The apparent predilection for events to occur in these regions rather than 3’ UTRs poses questions about the role of splicing in protein synthesis from translational regulation to the formation of polypeptide chains. Answering these questions requires a more precise understanding of how variation in UTRs and coding sequences affects final protein output as well as the biophysical characteristics of protein isoforms and their roles in different biological systems. These factors, combined with the large fraction of unannotated events in several cell types, should encourage tissue specialists to more deeply consider the contribution of isoform variation to cell identity and cell and tissue homeostasis.

Despite the clear association between splicing and cell identity, our analyses are yet to produce instances in which clustering in the splicing latent space reveals new cell subpopulations not visible in the expression latent space. This, of course, does not preclude the possibility in other settings where alternative splicing is known to be important, such as in specific developmental transitions or disease conditions. Nevertheless, our current experience leads us to believe that gene expression and splicing proportions provide two different projections of the same underlying cell state. Incidentally, RNA Velocity [45] estimates can be distorted by alternative splicing, and Bergen *et al*. [46] discuss incorporating isoform proportions into the model as a future direction.

To support our understanding of cell-type-specific splicing, we implemented a regularized generalized linear regression model which exploits the natural variation of splicing factors in different cell types. We recovered a number of previously identified (via knockout experiments) regulatory interactions and propose novel regulatory interactions involving genes known to play important regulatory roles. A key component of our analysis is the decision to include both the expression and alternative splicing patterns of splicing factors as features in the model. Consequently, we infer that several alternative splicing events in splicing factors themselves (some previously unannotated) contribute to their regulatory activity. Our model thus provides several opportunities for follow-up and does so with an increased granularity that distinguishes between effects due to expression and splicing differences. To facilitate further exploration of these data, we have uploaded our results to cell and genome browsers (linked at https://github.com/songlab-cal/scquint-analysis/).

Our experience analyzing these large data sets, initially with prior methods and then scQuint, has led to a series of general observations regarding the analysis of splicing in scRNA-seq data. As most analyses use full-length short-read protocols because of the cost of long-read data and the necessary focus on the 3’ end of transcripts in most UMI-based techniques, we restrict our attention to the full-length short-read setting and its incumbent challenges. For example, low transcript capture efficiency introduces additional technical noise into isoform quantification [47, 48, 49], and incomplete transcriptome annotations result in discarded reads and reduced sensitivity to cross-cell differences [48]. Nonetheless, we considered several methods (summarized in Table S1) to analyze isoform variation in short-read, full-length scRNA-seq. We found each of the classes of current methods to be problematic in the context of our data sets for varying reasons. Methods which depend on transcript annotations, [10, 50, 51, 52, 53, 54], cannot easily identify unannotated alternative splicing events. In large collections of previously unsurveyed cell types, these may comprise a sizable fraction of events. Indeed, we found up to 30% of differential splicing events were unannotated in certain cell types. Annotation-free approaches are also available, though possess their own limitations. For example, they may struggle to detect transcript variation involving small exons [12], or only consider binary alternative splicing events such as skipped or mutually exclusive exons [17, 18], ignoring other highly significant events such as alternative transcription start sites. Others propose *de novo* transcript assembly, but do not provide a statistical test for differential transcript usage across conditions [55, 23], limiting their immediate utility for exploration and interpretation of differences between cell types. Finally, methods can be divided into global or local quantifications based on the manner in which they featurize reads. Global methods, which examine the intron usage patterns across the entire gene, are greatly affected by coverage biases, of which we found many across both data sets, perhaps because so many cells and experiments comprise the atlases. These methods thus led to erroneous inference of cell clusters due to technical rather than biological variation. Conversely, local methods quantify splicing using events which occur near one another, reducing the effect of coverage artifacts along genes. Until the prevalence and severity of coverage biases are better understood, we advocate quantifying transcript variation in a local manner.

Recent and future experimental advances will catalyze the study of isoform variation in single cells. For instance, Smart-seq3 [56] allows sequencing of short reads from the entire length of a gene together with unique molecular identifiers, improving mRNA capture and allowing for the filtering of PCR duplicates; however, experiments show that less than 40% of reads can be unambiguously assigned to a single (annotated) isoform. Ultimately, long-read scRNA-seq will provide the definitive picture of isoform variation between cells. Until then, there is much biology to be studied using short-read protocols, and variation at the isoform level should not be disregarded.

## Materials and Methods

### Data sets

*Tabula Muris* data have accession code GSE109774. Cells were filtered to those from three month-old mice present in this collection: https://czb-tabula-muris-senis.s3-us-west-2.amazonaws.com/Data-objects/tabula-muris-senis-facs-processed-official-annotations. h5ad (filtering details in [57]). *BICCN Cortex* data were downloaded from https://assets.nemoarchive.org/dat-ch1nqb7 and filtered as in [22].

### Quantification

The bioinformatic pipeline was implemented using Snakemake [58]. Raw reads were trimmed from Smart-Seq2 adapters using Cutadapt [59] before mapping to the GRCm38/mm10 genome reference(https://hgdownload.soe.ucsc.edu/goldenPath/mm10/chromosomes/) and the transcriptome reference from Ensembl release 101 (ftp://ftp.ensembl.org/pub/release-101/gtf/mus_musculus/Mus_musculus.GRCm38.101.gtf.gz). Alignment was done using STAR [60] two-pass mode allowing novel junctions as long as they were supported by reads with at least 20 base pair overhang (30 if they are non-canonical) in at least 30 cells. Also, multimapping and duplicate reads were discarded using the flag `--bamRemoveDuplicatesType UniqueIdentical` (while this can remove duplicates from the second PCR step of Smart-seq, it will not remove duplicates from the first PCR step). Soft-clipped reads were removed as well. Additionally, reads were discarded if they belonged to the ENCODE region blacklist [61] (downloaded from https://github.com/Boyle-Lab/Blacklist/raw/master/lists/mm10-blacklist.v2.bed.gz).

Gene expression was quantified using featureCounts [62], and total-count normalized such that each cell had 10,000 reads (as in the Scanpy [63] tutorial). Intron usage was quantified using split reads with an overhang of at least 6 base pairs. Introns were discarded if observed in fewer than 30 cells in *BICCN Cortex* or 100 cells in *Tabula Muris*. Introns were grouped into alternative intron groups based on shared 3’ splice acceptor sites. Introns not belonging to any alternative intron group were discarded. Additionally, we decided to subset our analysis to introns with at least one of their donor or acceptor sites annotated, so we could assign a gene to them and facilitate interpretation for our specific analyses.

### Dimensionality reduction

To run PCA, we worked with alternative intron proportions (PSI) rather than their absolute counts, as the latter would be confounded by gene expression differences. However, given the sparsity of single-cell data, a very high proportion of alternative intron groups will have no reads in a given cell, leaving PSI undefined. More generally, an intron group may contain few reads, resulting in defined but noisy PSI estimates. To navigate this issue, we introduce a form of empirical shrinkage towards a central value. We first define the “global PSI” by aggregating reads from all cells and normalizing. Then, we add this global PSI as a pseudocount vector to each cell before re-normalizing to obtain each cell’s shrunken PSI profile (these are non-uniform pseudocounts adding up to one). We then run standard PCA on the cell-by-intron-smoothed PSI matrix.

The VAE was implemented using PyTorch [64] and scvi-tools [65]. The following is the generative model, repeated for each cell:

1. Sample the latent cell state *z* ~ Normal(0, I)
2. For each intron group *g*:

a. Obtain the underlying intron proportions:

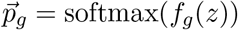
b. Sample the intron counts conditioning on the total observed *n_g_*:

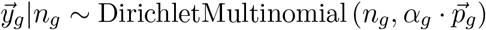

Here *f_g_*, known as the decoder, can be any differentiable function, including linear mappings and neural networks. *α_g_* is a scalar controlling the amount of dispersion. We optimize a variational posterior on cell latent variables *q*(*z*|*y*) (Gaussian with diagonal covariance, given by an encoder neural network) as well as point estimates of global parameters *f_g_*, *α_g_*. The encoder takes as input the smoothed PSI values, as in PCA, but the likelihood is based on the raw intron counts. The objective to maximize is the evidence lower bound (ELBO), consisting of a reconstruction term and a regularization term:

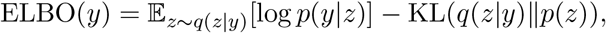

where KL(·||·) denotes the Kullback-Leibler divergence. Optimization is performed using Adam [66], a stochastic gradient descent method. To avoid overfitting in cases of relatively few cells with respect to the number of features, we considered a linear decoder [67], as well as a Normal(0, *σ*) prior on the entries of the decoder matrix. Hyperparameters were tuned using reconstruction error on held-out data and are described in Table S7.

To verify that we prevent leakage of gene expression information into our splicing profiles, we apply our VAE to embed a perturbed data set where intron counts within each intron group are redistributed with a fixed probability in all cells. This shuffled data set contains expression variability between cells but no splicing differences, and the splicing latent space does not distinguish among cell types (Figure S6).

### Differential splicing test

Our differential splicing test across conditions (such as cell types) is based on a Dirichlet-Multinomial Generalized Linear Model which was proposed in LeafCutter [16] for bulk RNA-seq. For each intron group *g* with *L* alternative introns:

- 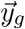 is a vector of counts for each of the *L* introns;
- The independent variable, *x*, equals 0 in one condition and 1 in the other;
- 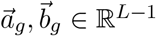 are the intercept and coefficients of the linear model;
- 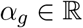 is a dispersion parameter shared across conditions; and
- the function softmax: 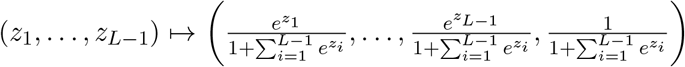 maps from 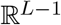 to the (*L* – 1)-dimensional probability simplex.

The Dirichlet-Multinomial Generalized Linear Model then proceeds as follows:

1. Obtain the underlying intron proportions:

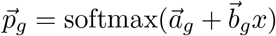
2. Sample the intron counts conditioned on the total observed, n*g*:

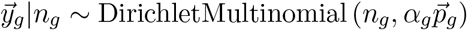

We implemented this model in PyTorch and optimized it using L-BFGS [68].

To test for differential splicing across the two conditions, we compare the following two hypotheses:

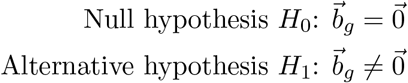

We use the likelihood-ratio test, the test statistic for which is asymptotically distributed as a χ^2^ random variable with *L* – 1 degrees of freedom under *H*_0_. Finally, we correct *p*-values for multiple testing using the Benjamini-Hochberg FDR procedure [69].

### Latent space analysis

The expression latent space was obtained by running PCA with 40 components on log-transformed and normalized gene expression values. The splicing latent space was obtained by running the VAE on the alternative intron count matrix (or equivalent features, e.g., Kallisto transcript counts, DEXSeq exon counts). Both latent spaces were visualized using UMAP [70]. In the comparison of Figure 1, we used our own implementation of the quantifications proposed by ODEGR-NMF, DEXSeq, and DESJ for ease of application to large single-cell datasets.

Dendrograms were constructed using hierarchical clustering (R function hclust) based on euclidean distance between the median latent space embedding of cells of each type. Tanglegram and entanglement were calculated using the dendextend R package, with the step2side method, as also described in [8].

Reported scores for cell type classification within a tissue were obtained by running a binary logistic regression classifier over 30 different random (stratified) splits with 2/3 of the data for training and 1/3 for testing.

### Cell-type-specific differential splicing

For differential splicing testing between a given cell type and the rest of the tissue, we only considered introns expressed in at least 50 cells and intron groups with at least 50 cells from both of the conditions. We called an intron group “differentially spliced” if it was both statistically significant using a 5% FDR and if it contained an intron with a PSI change greater than 0.05. We considered a differentially spliced intron group as unannotated if it contained an unannotated intron with a PSI change greater than 0.05. Differential expression was performed using the Mann-Whitney test. A gene was considered differentially expressed if it was statistically significant using a 5% FDR and if the fold change was at least 1.5.

### Splicing factor regression analysis

We obtained 75 mouse splicing factors using the Gene Ontology term “alternative mRNA splicing, via spliceosome” (http://amigo.geneontology.org/amigo/term/GO:0000380). A skipped exon annotation, processed by BRIE [51], was downloaded from https://sourceforge.net/projects/brie-rna/files/annotation/mouse/gencode.vM12/SE.most.gff3/download. Instead of using single cells as replicates, we partitioned the BICCN primary motor cortex dataset into roughly 200 clusters of 30 cells each that were pooled to create pseudobulks, aiming to reduce variance in the expression and splicing of splicing factors used as covariates in the model. We filtered target exon skipping events to those defined in at least 95% of the replicates, and those having a PSI standard deviation of at least 0.2. We used log-transformed normalized expression and PSI of alternative splicing events as input features. We chose to keep the PSI of only one intron per intron group to avoid the presence of highly correlated features and improve clarity, even if some information from non-binary events is lost. Input features were filtered to those having standard deviation of at least 0.05, and then standardized. A lasso Dirichlet-Multinomial GLM was fit to the data (in this instance, the model reduces to a Beta-Binomial because skipped exons are binary events), with the sparsity penalty selected via cross-validation. As a first approach, we fit a regular lasso linear regression model on PSI instead of raw counts, resulting in roughly similar patterns in the coefficients. Figure 10c shows the coefficients of the lasso Dirichlet-Multinomial model for the top 30 targets with the highest variance explained by the regular lasso model, all above 68%.

## Code and data availability

scQuint implementation in Python is available at https://github.com/songlab-cal/scquint. Differential splicing results and access to cell and genome browsers, together with code to reproduce results, are available at https://github.com/songlab-cal/scquint-analysis. Processed alternative intron count matrices are provided in the AnnData format (anndata.readthedocs.io) for easy manipulation with Scanpy [63], Seurat [71], and other tools.

## Acknowledgments

We would like to thank Angela Oliveira Pisco, Spyros Darmanis, and Kif Liakath-Ali for helpful discussions. We also thank the Chan Zuckerberg Biohub for hosting our cell×gene sessions and Aaron McGeever for assistance. This research is supported in part by grant number R35-GM134922 from NIH and grant number CZF2019-002449 from the Chan Zuckerberg Initiative Foundation. Y.S.S. is a Chan Zuckerberg Biohub Investigator.

