## Supplementary Information for "Robust and annotation-free analysis of alternative splicing across diverse cell types in mice"

This file contains:

- Supplementary Text
- Supplementary Tables S1–S7
- Supplementary Figures S1–S6

### Supplementary Text

#### Overview of available methods for alternative splicing analysis in full-length scRNA-seq data

Due to experimental considerations, the analysis of isoform variation in 10x Chromium data is mostly restricted to the 3' end of genes; in contrast, Smart-seq2 and other full-length, short-read protocols theoretically enable characterization of isoform variation along the whole gene. Nevertheless, numerous challenges impede such analyses in practice. For example, low transcript capture efficiency introduces additional technical noise into isoform quantification (Arzalluz-Luque and Conesa, 2018; Westoby et al., 2020; Buen Abad Najar et al., 2020), and incomplete transcriptome annotations result in discarded reads and reduced sensitivity to cross-cell differences (Westoby et al., 2020). Some authors have even recommended avoiding the analysis of alternative splicing in single-cell RNA sequencing (scRNA-seq) data until such obstacles can be suitably overcome (Westoby et al., 2020). Despite these difficulties, several methods (summarized in Table S1) have sought to analyze isoform variation in short-read, full-length scRNA-seq. Many methods, including kallisto (Bray et al., 2016), Census (Qiu et al., 2017), BRIE (Huang and Sanguinetti, 2017), SCATS (Hu et al., 2020), Quantas (Yan et al., 2015), VALERIE (meant only for visualization) (Wen et al., 2020) and BRIE2 (Huang and Sanguinetti, 2021), depend on transcript annotations and consequently cannot easily identify unannotated alternative splicing events, which may comprise a sizable fraction of events. Annotation-free approaches are also available, though they possess their own limitations. For example, ODEGR-NMF (Matsumoto et al., 2020) examines disparities in read coverage in fixed-size genomic windows but struggles to detect transcript variation involving small exons. Meanwhile, Expedition (Song et al., 2017) and ASCOT (Ling et al., 2020) only consider binary alternative splicing events such as skipped or mutually exclusive exons, ignoring other highly significant events such as alternate transcription start sites (TSS). SingleSplice (Welch et al., 2016) and RNA-Bloom (Nip et al., 2020) propose *de novo* transcript assembly, but do not provide a statistical test for differential transcript usage across conditions. DESJ (Liu et al., 2021) and SICILIAN/SpliZ (Dehghannasiri et al., 2021; Olivieri et al., 2021) compare intron proportions across the whole gene rather than locally, which we find is particularly sensitive to technical artifacts. Table S1 summarizes this information and makes the comparison of different methods easier.

#### Supplementary Tables

| Method | Isoform quantification | Annotation-free | Differential transcript usage |
|---|---|---|---|
| Quantas (Yan et al., 2015) | Local | | ✓ |
| SingleSplice (Welch et al., 2016) | Global | ✓ | * |
| kallisto (Bray et al., 2016) | Global | | * |
| Census (Qiu et al., 2017) | Global | | * |
| BRIE (Huang and Sanguinetti, 2017) | Local | | ✓ |
| Expedition (Song et al., 2017) | Local | ✓ | |
| ODEGR-NMF (Matsumoto et al., 2020) | Global | ✓ | * |
| SCATS (Hu et al., 2020) | Local | | ✓ |
| RNA-Bloom (Nip et al., 2020) | Global | ✓ | |
| ASCOT (Ling et al., 2020) | Local | ✓ | |
| DESJ (Liu et al., 2021) | Global | ✓ | ✓ |
| SpliZ (Olivieri et al., 2021) | Global | ✓ | ✓ |
| BRIE2 (Huang and Sanguinetti, 2021) | Local | | ✓ |
| **scQuint** | Local | ✓ | ✓ |

Table S1: **Summary of methods available to analyze transcript variation in short-read full-length scRNA-seq.** *Isoform quantification*: Does the method quantify global transcript variation or local alternative events? *Annotation-free*: Does quantification require an accurate transcriptome reference? *Differential transcript usage*: Does the method provide a two-sample test for differences in transcript proportions? Some methods, denoted by (*), provide other statistical tests. SingleSplice tests for alternative splicing within a single sample. kallisto and ODEGR-NMF test for differential transcript expression, i.e., changes in absolute transcript expression rather than their proportions. Census tests for differential transcript usage along a pseudotime trajectory.

| Data set | Cells | Cell types | Genes | Alt. introns | Unannotated |
|---|---|---|---|---|---|
| BICCN Cortex | 6220 | 11 | 26488 | 39357 | 29% |
| Tabula Muris | 44518 | 117 | 27348 | 29965 | 25% |

Table S2: **Overview of analyzed data sets.** Number of cells, cell types, detected genes, and detected alternative introns (including the percentage of introns that are not present in the Ensembl reference) for both data sources.

| Intron id | Intron coordinate |
|---|---|
| Ablim2_32870 | chr5:35828183-35833053 |
| Amz1_33732 | chr5:140724472-140741065 |
| Arhgap32_37706 | chr9:32208266-32239262 |
| Arhgef2_30803 | chr3:88616655-88629787 |
| Caly_16615 | chr7:140081539-140082462 |
| Camk2a_28025 | chr18:60963964-60969994 |
| Cpeb1_15988 | chr7:81372201-81372409 |
| Dlc1_16933 | chr8:36599441-36763645 |
| Dlgap1_27522 | chr17:70516997-70593167 |
| Etv1_23259 | chr12:38861106-38861287 |
| Faap20_32501 | chr4:155246986-155250506 |
| Fhit_4932 | chr14:10421614-10453369 |
| Frmd4b_14813 | chr6:97423525-97487593 |
| Frmpd4_19441 | chrX:167897639-168447948 |
| Gas7_22076 | chr11:67455775-67603108 |
| Gria1_21954 | chr11:57310723-57317666 |
| Gria2_11163 | chr3:80689352-80690404 |
| Grip1_21379 | chr10:119692622-119897715 |
| Grm5_35660 | chr7:88075131-88129981 |
| Kalrn_6826 | chr16:34055116-34095796 |
| Macf1_12504 | chr4:123440792-123441603 |
| Nek11_18537 | chr9:105348139-105348542 |
| Nrg1_16909 | chr8:31849440-31917537 |
| Nrxn1_8067 | chr17:90597630-90623420 |
| Oxr1_25579 | chr15:41831130-41848670 |
| Pgm2_32951 | chr5:64095936-64096979 |
| Ppfia2_21330 | chr10:106632879-106743487 |
| Pstpip2_28237 | chr18:77794751-77835125 |
| Ptk2_5956 | chr15:73282236-73283267 |
| Ptprd_12131 | chr4:76136922-76138556 |
| Rapgef4_29435 | chr2:72174477-72174802 |
| Rbfox1_26172 | chr16:5763912-6173605 |
| Rcan2_27335 | chr17:43836498-44017767 |
| Shisa9_26238 | chr16:12244764-12267424 |
| Slmap_5079 | chr14:26414989-26418113 |
| Sox5_15181 | chr6:144155224-144434425 |
| Spats1_7721 | chr17:45464581-45539630 |
| Srr_2782 | chr11:74913134-74925619 |
| Tiam1_7094 | chr16:89813128-89817961 |
| Vgll4_14901 | chr6:114890807-114921317 |

Table S3: Intron coordinates for Figure 5.

| Intron id | Intron coordinate |
|---|---|
| Capg_26086 | chr6:72544467-72555273 |
| Dock10_327 | chr1:80648333-80758326 |
| Foxp1_11076 | chr6:99260420-99266347 |
| Gsn_22212 | chr2:35266904-35283905 |
| Rps6ka1_9389 | chr4:133872004-133874608 |
| Selplg_10200 | chr5:113820249-113832429 |
| Smarca4_28720 | chr9:21626013-21632805 |
| Syk_18319 | chr13:52597055-52610796 |

Table S4: Intron coordinates for Figure 7.

| Intron id | Intron coordinate |
|---|---|
| Cela1_4832 | chr15:100681289-100681794 |
| Entpd5_3004 | chr12:84399413-84405406 |
| Eps8_11340 | chr6:137539402-137570702 |
| Fam107b_21905 | chr2:3713637-3772796 |
| Fam110a_7811 | chr2:151970925-151979993 |
| Fermt3_6600 | chr19:6999781-7001757 |
| Fxyd2_28841 | chr9:45406167-45408106 |
| Ggt1_15810 | chr10:75564906-75573611 |
| Gsn_22213 | chr2:35282602-35283905 |
| Itpr1_26257 | chr6:108493894-108499108 |
| Khk_24896 | chr5:30927157-30928443 |
| Limk2_1649 | chr11:3360587-3370969 |
| Mapre2_21026 | chr18:23772660-23832853 |
| Nedd4l_21245 | chr18:64997422-65074724 |
| Pla2g4a_577 | chr1:149922137-149922244 |
| Plac8_10073 | chr5:100562763-100562841 |
| Prkacb_8667 | chr3:146757769-146781147 |
| Prlr_19174 | chr15:10216124-10293651 |
| Rbm47_9908 | chr5:66046214-66097707 |
| Sectm1a_2756 | chr11:121075722-121081083 |
| Tln2_13865 | chr9:67459990-67485118 |
| Tprg_19889 | chr16:25385997-25412767 |
| Vps29_25440 | chr5:122356806-122360013 |
| Akap13_27114 | chr7:75643465-75663412 |
| Aplp2_13526 | chr9:31153500-31157695 |
| App_5359 | chr16:85030366-85040251 |
| Arid5a_14716 | chr1:36317663-36318084 |
| C1qtnf1_17475 | chr11:118428569-118443692 |
| Egfl7_22064 | chr2:26581242-26585585 |
| Emcn_23704 | chr3:137391707-137416721 |
| Fhl1_29631 | chrX:56786583-56787989 |
| Hyal2_29252 | chr9:107567977-107570104 |
| Lims1_15716 | chr10:58394601-58399021 |
| Lmo2_22492 | chr2:103970366-103970493 |
| Myl6_1604 | chr10:128491034-128492059 |
| Ndufa7_20560 | chr17:33825651-33829667 |
| Pecam1_2522 | chr11:106679612-106682880 |
| Pkm_28967 | chr9:59675196-59678044 |
| Pllp_13202 | chr8:94677349-94679345 |
| Pparg_26324 | chr6:115362418-115383679 |
| Rbms1_7374 | chr2:60752983-60755416 |
| Rps24_18700 | chr14:24493494-24495775 |
| S100a16_23418 | chr3:90537418-90541994 |
| Slc39a10_137 | chr1:46836152-46892800 |
| Slc9a3r2_5546 | chr17:24642371-24644819 |
| Slfn5_16990 | chr11:82952196-82956263 |
| Spry1_23161 | chr3:37640809-37642555 |
| Srsf2_2675 | chr11:116850962-116851648 |
| Srsf7_6089 | chr17:80204388-80204622 |
| St3gal6_5296 | chr16:58507615-58523502 |
| Tacc1_12778 | chr8:25177995-25240794 |
| Tcf4_21297 | chr18:69564685-69633577 |
| Tmcc3_16030 | chr10:94515105-94578423 |
| Txnrd1_15967 | chr10:82859301-82874613 |
| Ube2d3_23688 | chr3:135438699-135439644 |

Table S5: Intron coordinates for Figure 8.

| Intron id | Intron coordinate |
|---|---|
| Ablim2_32871 | chr5:35831743-35833053 |
| Ablim2_32873 | chr5:35839823-35841281 |
| Ablim2_32875 | chr5:35843267-35848804 |
| Ahi1_20693 | chr10:21070544-21072543 |
| Cadps_4944 | chr14:12467180-12468346 |
| Camk2a_28026 | chr18:60969044-60969994 |
| Camta1_12911 | chr4:151062856-151071426 |
| Celf2_9334 | chr2:6604183-6607882 |
| Celf2_9349 | chr2:6721614-6962379 |
| Cttn_16755 | chr7:144449616-144450978 |
| Dlg2_35711 | chr7:91997322-92040798 |
| Dlgap1_27532 | chr17:70718226-70761075 |
| Fam172a_24454 | chr13:77761939-77825323 |
| Fxr1_30525 | chr3:34064320-34068161 |
| Gabrg2_2316 | chr11:41912592-41913970 |
| Golga1_9791 | chr2:39052102-39052956 |
| Gria2_11163 | chr3:80689352-80690404 |
| Grin1_9480 | chr2:25310476-25311343 |
| Hnrnpa1_26117 | chr15:103243081-103243625 |
| Hnrnpm_7568 | chr17:33669177-33677274 |
| Ints10_36756 | chr8:68824268-68825052 |
| Kcnma1_5069 | chr14:23442080-23449200 |
| Khdrbs3_25689 | chr15:68929658-68995052 |
| Kif21a_6295 | chr15:90972716-90976471 |
| Mbnl1_30665 | chr3:60501424-60528756 |
| Mbnl2_25375 | chr14:120395766-120396451 |
| Mbnl2_25376 | chr14:120396605-120404638 |
| Mbnl2_25378 | chr14:120396605-120428935 |
| Nptn_37975 | chr9:58619011-58623687 |
| Nrxn1_8060 | chr17:90162434-90163792 |
| Nrxn1_8066 | chr17:90597630-90620865 |
| Nrxn2_28410 | chr19:6513806-6516923 |
| Nrxn3_23721 | chr12:90199237-90204511 |
| Phactr1_24192 | chr13:43059778-43077655 |
| Pqbp1_18852 | chrX:7898637-7899169 |
| Rap1gap_32329 | chr4:137723824-137724692 |
| Rapgef4_29435 | chr2:72174477-72174802 |
| Rbfox1_26172 | chr16:5763912-6173605 |
| Rbfox1_26177 | chr16:6173668-6487819 |
| Rbfox1_26185 | chr16:6809249-7224311 |
| Rbfox1_26203 | chr16:7392041-7407964 |
| Rbfox2_6060 | chr15:77086156-77094451 |
| Rbfox2_6066 | chr15:77132942-77231387 |
| Rbm25_23574 | chr12:83661269-83663979 |
| Rnps1_26974 | chr17:24415163-24418038 |
| Slmap_5078 | chr14:26414989-26416203 |
| Snap91_18411 | chr9:86792737-86793357 |
| Srsf9_33347 | chr5:115328705-115330498 |
| Srsf9_33351 | chr5:115331540-115332117 |
| Stxbp1_9672 | chr2:32789638-32794588 |
| Tex9_18291 | chr9:72478423-72479924 |
| Wdr17_16993 | chr8:54690235-54693057 |

Table S6: Intron coordinates for Figure 10 and Figure S5.

| Data set | Decoder | Layers | $\sigma$ | Latent dimension |
|---|---|---|---|---|
| BICCN Cortex | Linear | 1 | 26.8 | 18 |
| Tabula Muris | Non-linear | 2 | - | 34 |

Table S7: **VAE hyperparameters.**

### Supplementary Figures

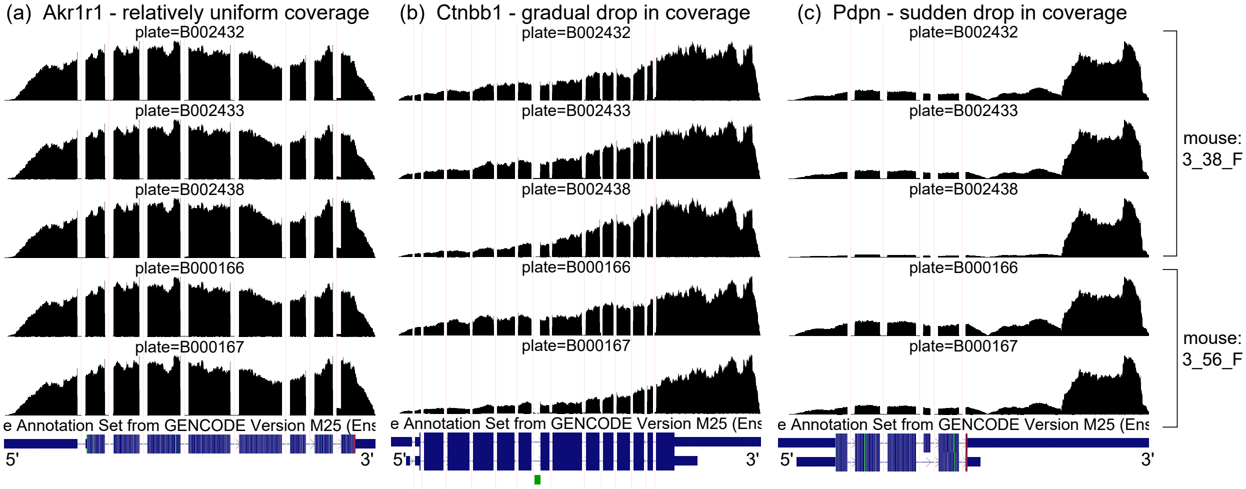


Figure S1: **Coverage artifacts in mammary gland basal cells from *Tabula Muris*.** Aggregate read coverage of basal cells is shown for three genes in two female mice: *3_38_F*, processed in three different plates, and *3_56_F*, processed in two different plates. Visualization on the UCSC Genome Browser. **(a)** *Akr1r1*, with relatively uniform coverage, what we expect. **(b)** *Ctnbb1*, with a gradual drop in coverage away from the 3' end. The rate of coverage decay varies across plates. **(c)** *Pdpn*, with a sudden drop in coverage halfway through the 3' UTR. The magnitude of the drop varies across plates.

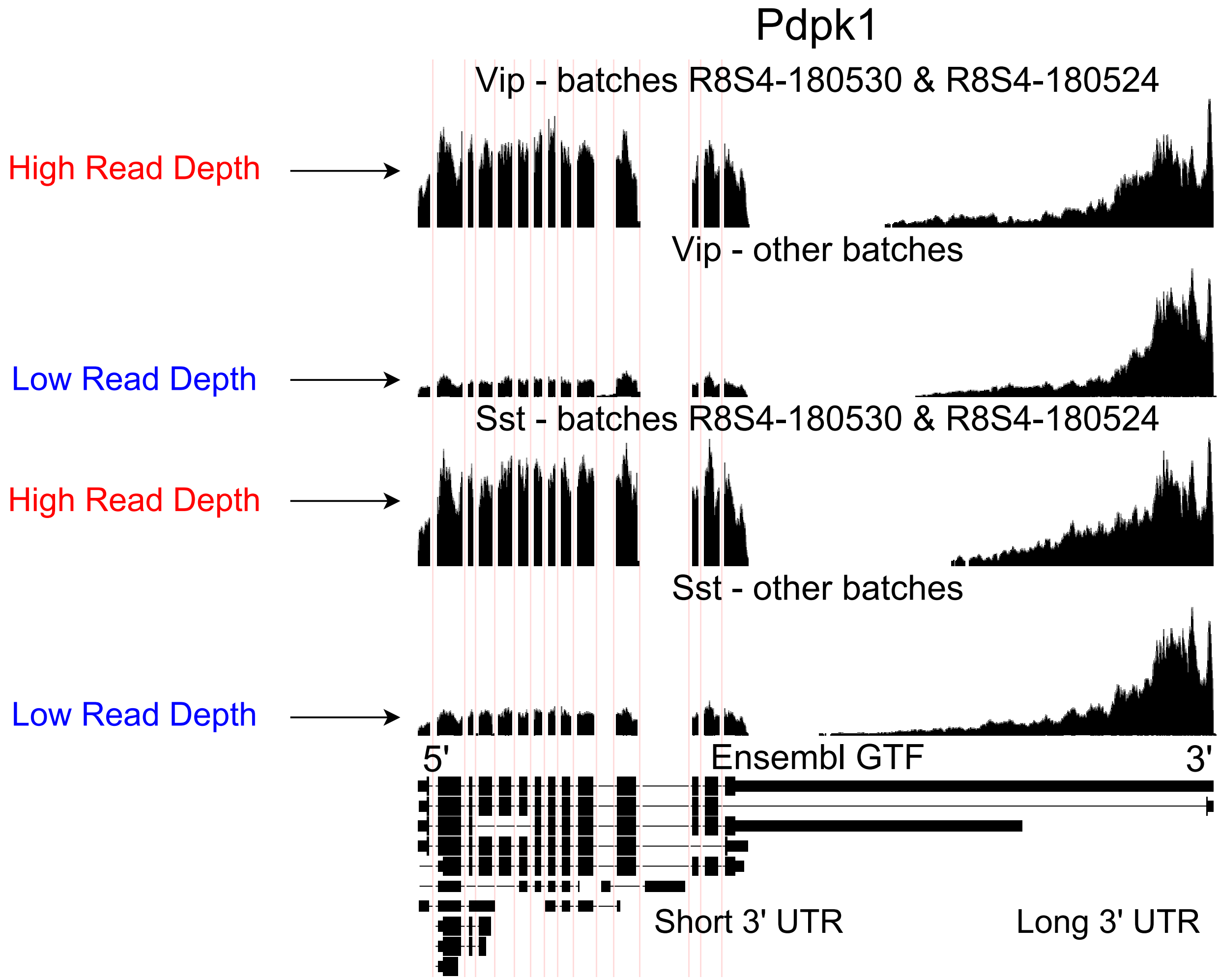


Figure S2: **Technical artifacts in *BICCN Cortex*.** Aggregate read coverage in *Pdpk1* in two cell types, Vip and Sst, further separated according to the "batch" metadata label into two groups. The first group contains cells from batches *R8S4-180530* and *R8S4-180524*, while the second group contains the remaining batches. Cells from different batches belong to different mice and were processed on different dates. In all groups, coverage decreases rapidly in the 3' UTR of the isoform with the longest 3' UTR, eventually reaching zero. Additionally, the relative coverage in this region compared to the rest of the gene (seeming to originate from a different isoform with a short 3' UTR) varies drastically across batches, and consistently in different cell types. In principle, this could be due to biological differences between mice from different batches, but an explanation based on technical factors such as amplification bias may be more plausible.

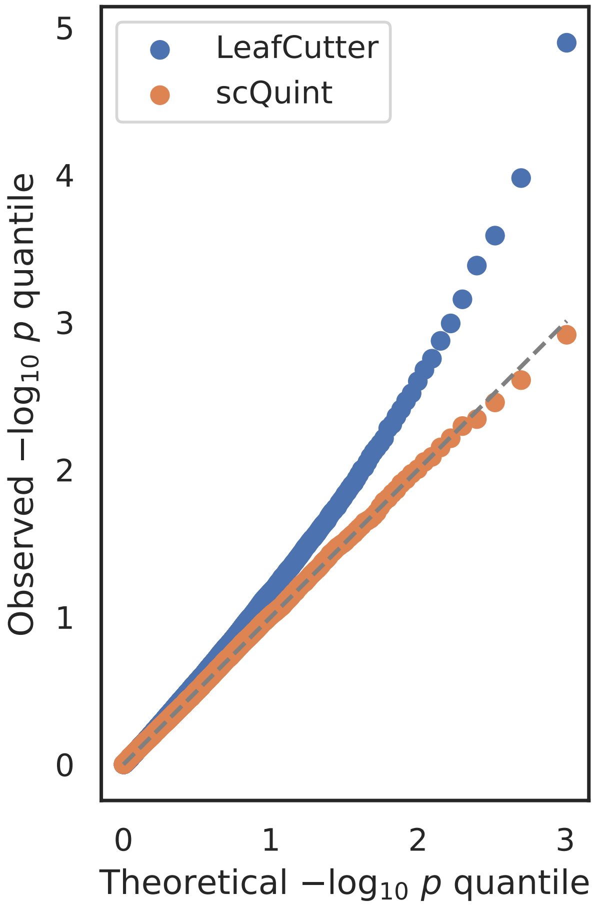


Figure S3: **Distribution of $p$-values under the null.** We randomly split 6220 *BICCN Cortex* cells into two equally sized groups and performed our test of differential splicing. In this scenario, the null hypothesis of no difference in splicing proportions holds, and we expect the distribution of $p$-values to be uniform. The quantile-quantile plot of $p$-values obtained with scQuint shows their distribution is indeed uniform, suggesting that the model is well-calibrated under the null; this is not true for $p$-values obtained by LeafCutter.

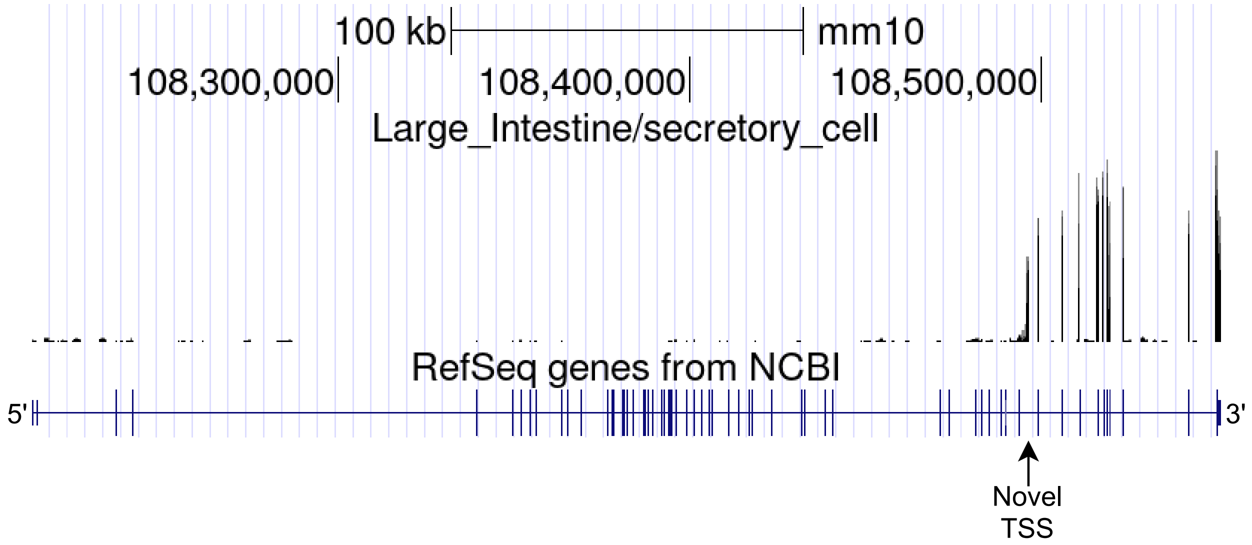


Figure S4: **Full-gene view of novel alternative TSS in *Itpr1*.** Large intestine secretory cells aggregate read coverage visualized in the UCSC Genome Browser.

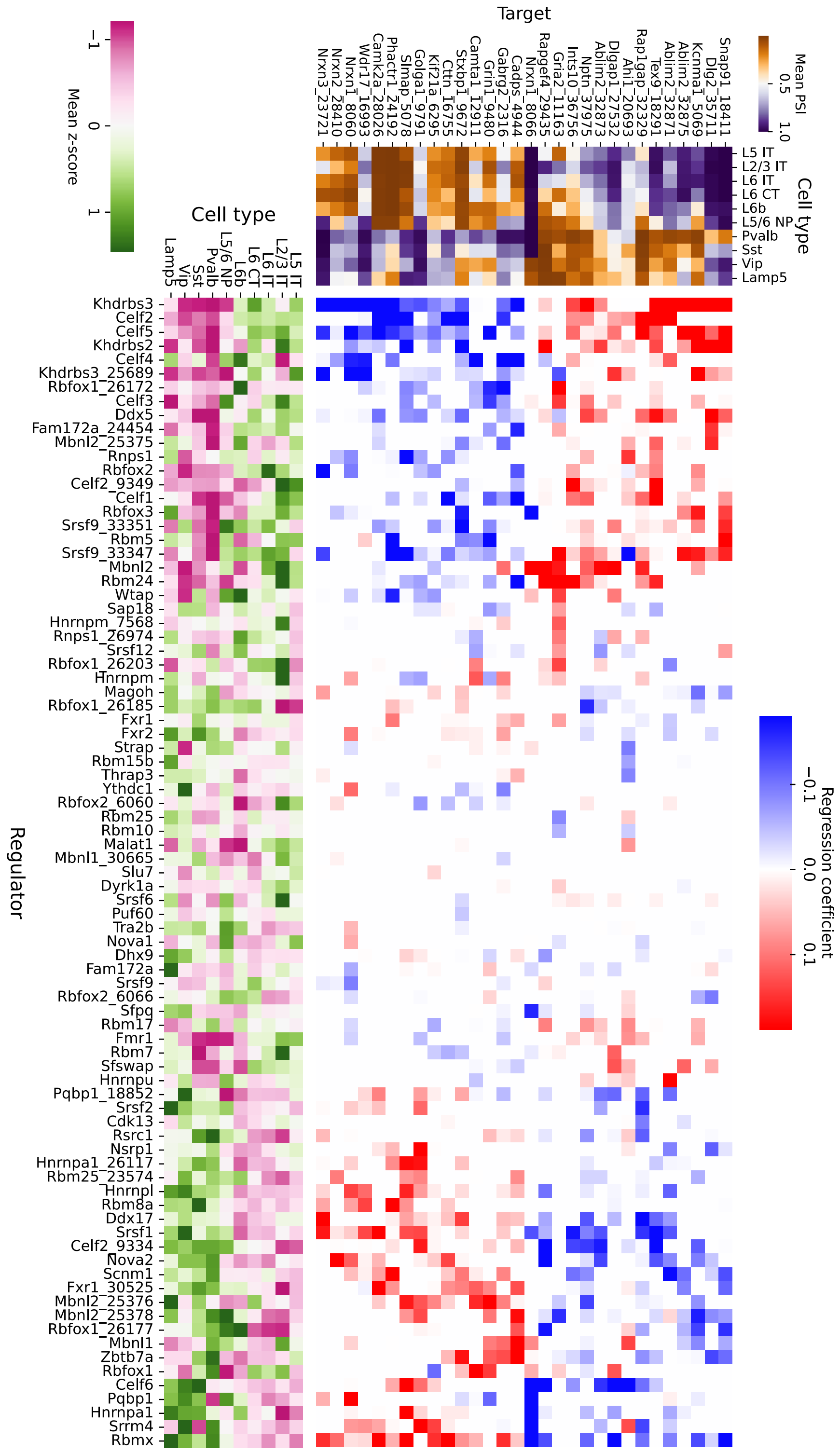


Figure S5: **Associations between splicing factors and alternative splicing (extended).** Regression analysis of exon skipping based on expression and splicing of splicing factors, using the BICCN mouse primary motor cortex dataset. Left panel: mean PSI of skipped exons across cell types. Bottom panel: mean z-scores of selected splicing factor features across cell types, including whole-gene expression (gene name) and PSI of alternative introns (gene name and numerical identifier). Intron coordinates are available in Table S6. Center panel: regression coefficients (log-odds) of each splicing factor feature used to predict skipped exon PSI in our sparse Dirichlet-Multinomial linear model.

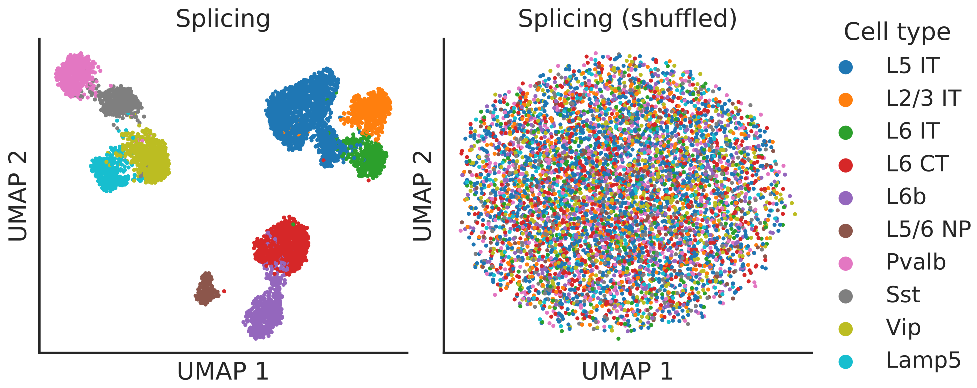


Figure S6: **Splicing latent space when alternative intron counts are shuffled.** To verify that absolute gene expression does not affect the splicing latent space, we perturbed the *BICCN Cortex* data set by resampling alternative intron counts with a fixed proportion in all cells (the proportions in different alternative intron groups varied and were sampled from a uniform Dirichlet distribution). In this scenario, different cell types still vary in their gene expression levels but not in their splicing patterns. As hoped, the splicing latent space does not distinguish between cell types, indicating it is only capturing differences in splicing proportions rather than changes in absolute gene expression.
